# Functional geometry of auditory cortical resting state networks derived from intracranial electrophysiology

**DOI:** 10.1101/2022.02.06.479292

**Authors:** Matthew I. Banks, Bryan M. Krause, D. Graham Berger, Declan I. Campbell, Aaron D. Boes, Joel E. Bruss, Christopher K. Kovach, Hiroto Kawasaki, Mitchell Steinschneider, Kirill V. Nourski

**Affiliations:** Department of Anesthesiology, University of Wisconsin, Madison, WI, USA; Department of Neuroscience, University of Wisconsin, Madison, WI, USA; Department of Neurology, The University of Iowa, Iowa City, IA 52242, USA; Department of Neurosurgery, The University of Iowa, Iowa City, IA 52242, USA; Department of Neurology, Albert Einstein College of Medicine, New York, NY 10461, USA; Department of Neuroscience, Albert Einstein College of Medicine, New York, NY 10461, USA; Iowa Neuroscience Institute, The University of Iowa, Iowa City, IA 52242, USA

**Keywords:** Functional connectivity, hierarchy, fMRI, diffusion map embedding, electrocorticography, intracranial electroencephalography

## Abstract

Understanding central auditory processing critically depends on defining underlying auditory cortical networks and their relationship to the rest of the brain. We addressed these questions using resting state functional connectivity derived from human intracranial electroencephalography. Mapping recording sites into a low-dimensional space where proximity represents functional similarity revealed a hierarchical organization. At fine scale, a group of auditory cortical regions excluded several higher order auditory areas and segregated maximally from prefrontal cortex. On mesoscale, the proximity of limbic structures to auditory cortex suggested a limbic stream that parallels the classically described ventral and dorsal auditory processing streams. Identities of global hubs in anterior temporal and cingulate cortex depended on frequency band, consistent with diverse roles in semantic and cognitive processing. On a macro scale, observed hemispheric asymmetries were not specific for speech and language networks. This approach can be applied to multivariate brain data with respect to development, behavior, and disorders.

**Blurb:** We describe the organization of human neocortex on multiple spatial scalesbased on resting state intracranial electrophysiology. We focus on cortical regions involved in auditory processing and examine inter-regional hierarchical relationships, network topology, and hemispheric lateralization. This work introduces a powerful analytical tool to examine mechanisms of altered arousal states, brain development, and neuropsychiatric disorders.

## Introduction

The meso- and macroscopic organization of human neocortex has been investigated extensively using resting state (RS) functional connectivity, primarily using functional magnetic resonance imaging (fMRI)[1, 2]. RS data are advantageous as they avoid the substantial confound of stimulus-driven correlations yet identify networks that overlap with those obtained using event-related data[3], and thus are relevant to cognitive and perceptual processing. RS fMRI has contributed greatly to our understanding of the organization of the human auditory cortical hierarchy[4–6], but only a few complementary studies have been conducted using electrophysiology in humans (e.g., Refs.[7–9]). Compared to fMRI, intracranial electroencephalography (iEEG) offers superior spatio-temporal resolution, allowing for analyses that accommodate frequency-dependent features of information exchange in these networks[10, 11]. For example, cortico-cortical feedforward versus feedback information exchange occurs via band-specific communication channels (gamma band and beta/alpha bands, respectively) in both the visual[11–15] and auditory[16–19] systems. There are also important regions involved in speech and language processing for which iEEG can provide superior spatial resolution and signal characteristics compared to fMRI, including in the anterior temporal lobe[20, 21] and the upper versus lower banks of the superior temporal sulcus (STS)[22, 23]. However, variable electrode coverage in human intracranial patients and small sample sizes are challenges to generalizing results.

We overcome these limitations using a large cohort of subjects that together have coverage over most of the cerebral cortex and leverage these data to address outstanding questions about auditory networks. We address the organization of human auditory cortex at three spatial scales: fine-scale organization of regions adjacent to canonical auditory cortex, clustering of cortical regions into functional processing streams, and hemispheric (a)symmetry associated with language dominance. We present a unified analytical framework applied to resting state human iEEG data that embeds functional connectivity data into a Euclidean space in which proximity represents functional similarity. A similar analysis has been applied previously to RS fMRI data[24–26]. We extend this analytical approach and demonstrate methodology appropriate for hypothesis testing at each of these spatial scales.

At the fine scale, individual areas within canonical auditory cortex and beyond have different sensitivity and specificity of responses with respect to stimulus attributes[27–29]. These differences are related to underlying connectivity patterns both within the auditory cortex and with other brain areas[22]. Though there is broad agreement that posteromedial Heschl’s gyrus (HGPM) represents core auditory cortex, functional relationships among HGPM and neighboring higher-order areas are still a matter of debate. For example, the anterior portion of the superior temporal gyrus (STGA) and planum polare (PP) are adjacent to auditory cortex on Heschl’s gyrus, yet diverge from it functionally[30, 31]. The posterior insula (InsP), on the other hand, has response properties similar to HGPM, yet is not considered a canonical auditory area[32]. The STS is a critical node in speech and language networks[22, 33–37], yet its functional relationships with other auditory areas are difficult to distinguish with neuroimaging methods. Indeed, distinct functional roles of its upper and lower banks (STSU, STSL) have only been recently elucidated with iEEG[23].

Questions remain regarding mesoscale organization as well. The auditory hierarchy is posited to be organized along two processing streams (ventral “what” and dorsal “where/audiomotor”) [38–40]. The specific brain regions involved and the functional relationships within each stream are vigorously debated[41–44]. Furthermore, communication between auditory cortex and hippocampus, amygdala, and anterior insula (InsA)[45] – areas involved in auditory working memory and processing of emotional aspects of auditory information[46–49] – suggests a third “limbic” auditory processing stream, complementary to the dorsal and ventral streams.

At a macroscopic scale, hemispheric lateralization is a classically described organizational feature of speech and language function[50, 51]. However, previous studies have shown extensive bilateral activation during speech and language processing[52–54], and more recent models emphasize this bilateral organization[39]. Thus, the degree to which lateralization shapes the auditory hierarchy and is reflected in hemisphere-specific connectivity profiles is unknown[38, 42, 55–58].

To address these questions, we applied diffusion map embedding (DME)[59, 60] to functional connectivity measured between cortical regions of interest (ROIs). DME is part of a broader class of analytical approaches that leverage the spectral properties of similarity matrices to reveal the intrinsic structure of datasets[61]. When applied to multivariate neurophysiological signals, DME maps connectivity from anatomical space (i.e., the location of the recording sites in the brain) into a Euclidean embedding space that reveals a “functional geometry”[24]. In this space, the proximity of two ROIs reflects similarity in connectivity to the rest of the network. Implicit in the use of the term ‘functional’ is the assumption that two regions of interest that are similarly connected to the rest of the brain are performing similar functions. Here, we use the DME approach to provide a low-dimensional representation convenient for display while also facilitating quantitative comparisons on multiple spatial scales. We tested pre-specified hypotheses of specific ROI relationships involving STSL and STSU in the gamma band using permutation tests. We applied exploratory analyses to other bands, to hierarchical clustering to identify functional processing streams, and to contrasts of whole embeddings between participant cohorts to investigate hemispheric differences in network organization.

This is the first time to our knowledge DME analysis has been applied to electrophysiological data, which allows exploration of the band-specificity of network structure. Also novel in our approach is the examination of relationships based on inter-ROI distances in embedding space, which are robust to changes in the underlying basis functions of the space.

## Results

### DME applied to iEEG data

Intracranial electrodes densely sampled cortical structures involved in auditory processing in the temporal and parietal lobes, as well as prefrontal, sensorimotor, and other ROIs in 49 participants (22 female; Fig. 1, Supplementary Tables 1, 2). A total of 6742 recording sites (66.1% subdural, 33.9% depth) were used in the analyses. On average, each participant contributed 138±54 recording sites, representing 28±7.7 ROIs (mean ± standard deviation) (see example in Fig. 2a). Figure 1b summarizes both subdural and depth electrode coverage by plotting recording sites in Montreal Neurological Institute (MNI) coordinate space and projecting them onto an average template brain for spatial reference. Of note, assignment of recording sites to ROIs as depicted in Figure 1 was made based on the sites’ locations in each participant’s brain rather than based on the projection onto the template brain, thus accounting for the high individual variability in cortical anatomy (see Methods for details).

**Figure 1.**
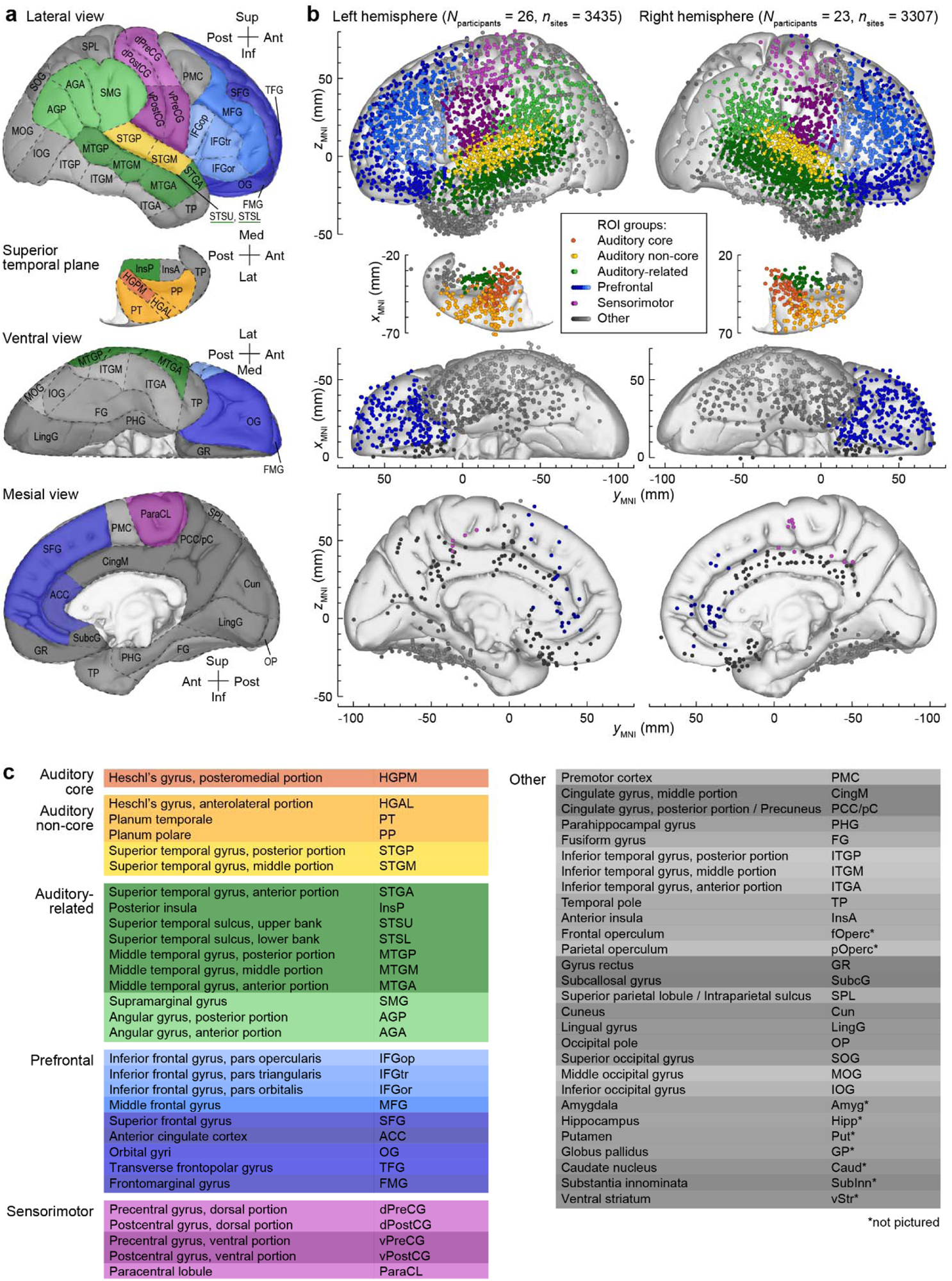
ROIs and electrode coverage in all 49 participants. **a:** ROI parcellation scheme. **b:** Locations of recording sites, determined for each participant individually and color-coded according to the ROI group, are plotted in Montreal Neurological Institute (MNI) coordinate space and projected onto the Freesurfer average template brain for spatial reference. Color shades represent different ROIs within a group. Projections are shown on the lateral, top-down (superior temporal plane), ventral and mesial views (top to bottom). Recording sites over orbital, transverse frontopolar, inferior temporal gyrus and temporal pole are shown in both the lateral and the ventral view. Sites in fusiform, lingual, parahippocampal gyrus and gyrus rectus are shown in both the ventral and medial view. Sites in the frontal operculum (*n* = 23), parietal operculum (*n* = 21), amygdala (*n* = 80), hippocampus (*n* = 86), putamen (*n* = 15), globus pallidus (*n* = 1), caudate nucleus (*n* = 10), substantia innominata (*n* = 5), and ventral striatum (*n* = 2) are not shown. See Supplementary Table 2 for detailed information on electrode coverage. **c:** ROI groups, ROIs and abbreviations used in the present study. See Supplementary Table 3 for alphabetized list of abbreviations.

**Figure 2.**
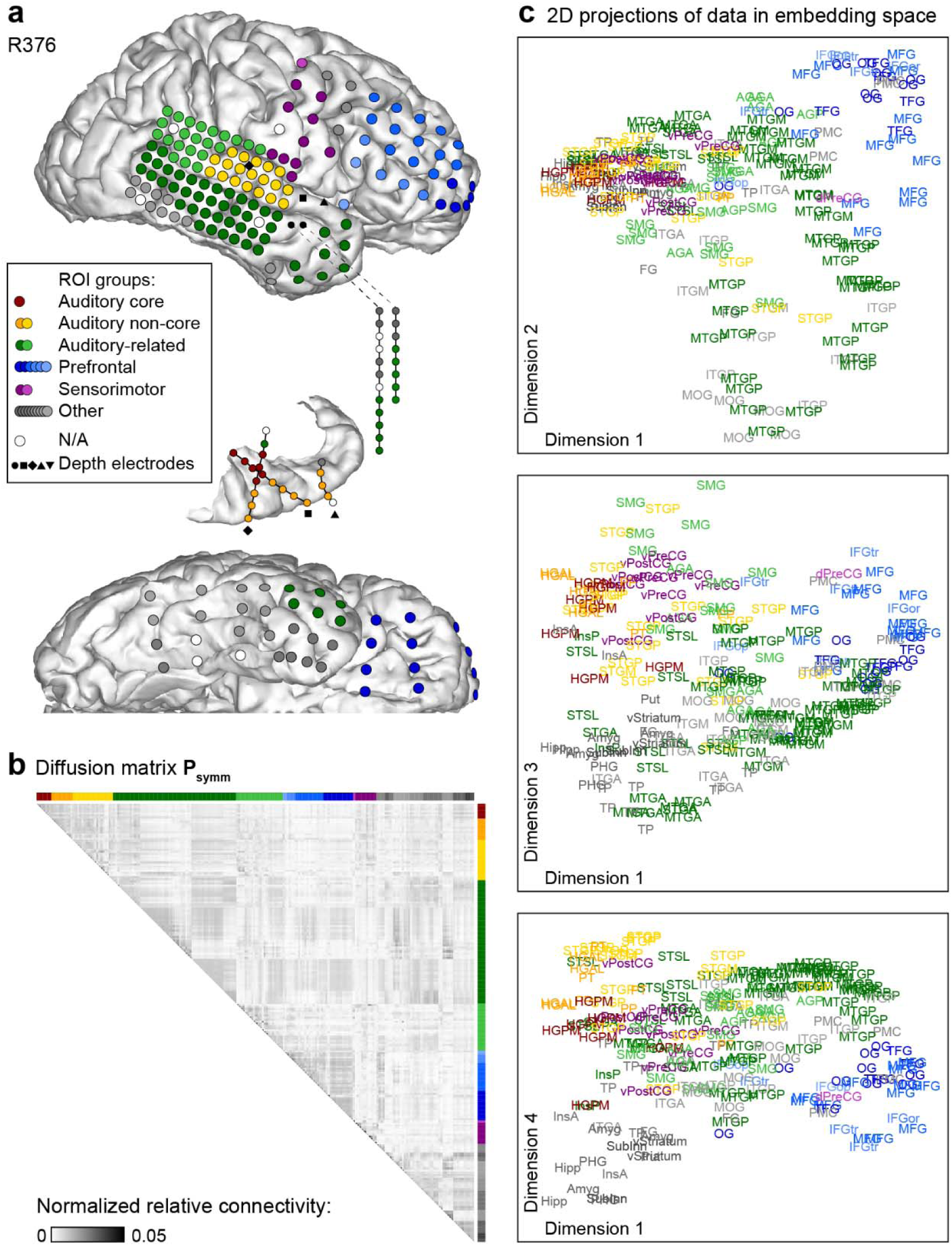
Functional geometry of cortical networks revealed by DME applied to gamma band power envelope correlations in a single participant (R376). **a:** Electrode coverage. **b:** Diffusion matrix **P_symm_**. **c:** Data plotted on the same scale in the 1st and 2nd, 1st and 3rd, and 1st and 4th dimensions of embedding space (top to bottom). Two points that are close in embedding space are similarly connected to the rest of the network, and thus assumed to be functionally similar.

The brain parcellation scheme depicted in Figure 1a was developed based on a combination of physiological and anatomical criteria and has been useful in our previous analyses that were largely focused on auditory processing[62–67]. One goal of the analysis presented in this study is to develop instead a parcellation scheme based on functional relationships between brain areas. Accordingly, we revisit below the parcellation shown in Fig. 1a with a data-driven scheme.

DME was applied to pairwise functional connectivity measured as orthogonalized power envelope correlations[68] computed between recording sites in each participant. We focus on gamma band power envelope correlations because of its established role in feedforward information exchange in the auditory system[16–19], and use gamma band as a reference in presentation of data from other bands. The functional connectivity matrix was normalized and thresholded to yield a diffusion matrix **P_symm_** with an apparent community structure along the horizontal and vertical dimensions (Fig. 2b). DME reveals the functional geometry of the sampled cortical sites by using the structure of **P_symm_** and a free parameter *t* to map the recording sites into an embedding space. In this space, proximity between nodes represents similarity in their connectivity to the rest of the network (Fig. 2c; see Supplementary Fig. 1 for additional views). The parameter *t* corresponds to diffusion time: larger values of *t* shift focus from local towards global organization. DME exhibited superior signal-to noise characteristics compared to direct analysis of functional connectivity in 43 out of 49 participants (Supplementary Fig. 2).

Functionally distinct regions are isolated along principal dimensions in embedding space. For example, in Figure 2c, auditory cortical sites (red/orange/yellow) and sites in prefrontal cortex (blue) were maximally segregated along dimension 1 (see Fig. 1 and Supplementary Table 3 for the list of abbreviations). Other regions (e.g., middle temporal gyrus) had a more distributed representation within the embedding space, consistent with their functional heterogeneity.

### Functional geometry of cortical networks

To pool data across participants with variable electrode coverage, **P_symm_** matrices were computed at the ROI level and averaged across participants. The results for gamma band data are shown in Figure 3a. The eigenvalue spectrum |λ*_i_*| of this averaged **P_symm_** showed a clear separation between the first four and the remaining dimensions (Fig. 3a, inset), indicating that the first four dimensions of embedding space accounted for much of the community structure of the data. Indeed, these first four dimensions accounted for >80% of the diffusion distance averaged across all pairwise distances in the space, a typical measure for deciding which dimensions to retain when DME is used as a dimensionality reduction method[60]. This inflection point in the eigenvalue spectrum was identified algorithmically (see Methods) for each frequency band and yielded the number of retained dimensions *n* = 6,6,7,4, and 6 for theta, alpha, beta, gamma, and high gamma bands, respectively.

**Figure 3.**
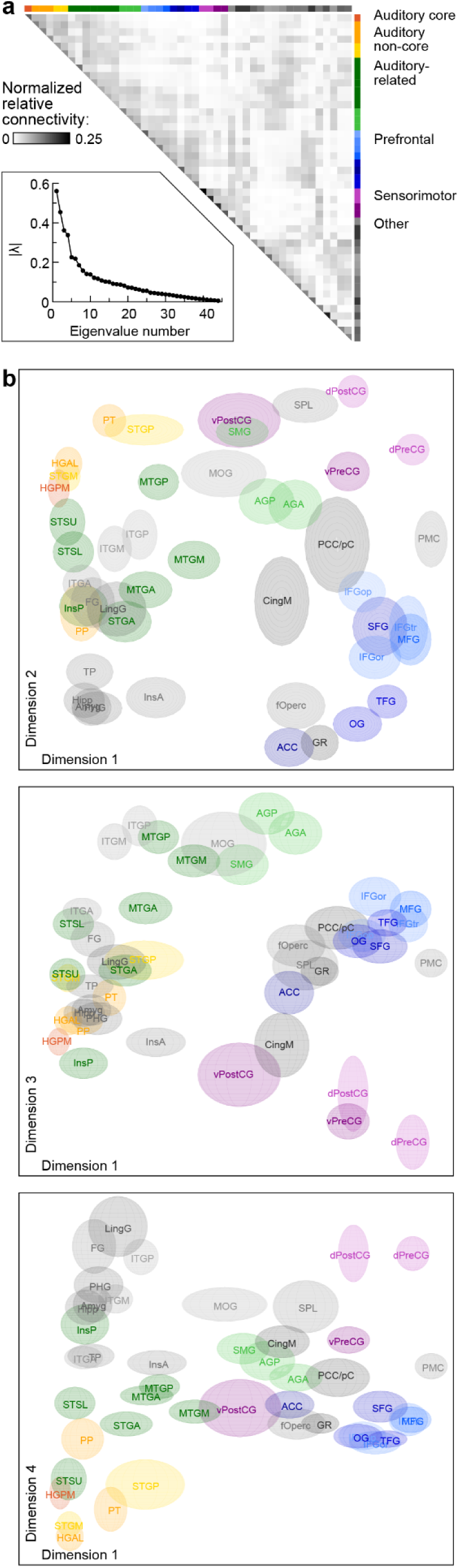
Summary of functional geometry of cortical networks via DME applied to gamma band power envelope correlations. **a**: Average diffusion matrix. **Inset:** Eigenvalue spectrum. **b:** Data plotted on the same scale in the 1st and 2nd, 1st and 3rd, and 1st and 4th dimensions of embedding space (top to bottom). Estimates of variance across participants in the locations of each ROI in embedding space were obtained via bootstrapping and are represented by the size of the ellipsoid for each ROI.

The gamma band data are plotted in the first four dimensions of embedding space in Figure 3b, where the sizes of the ellipsoids for each ROI represent estimates of position variance across participants obtained via bootstrapping. These data provide a graphical representation of the functional geometry of all sampled brain regions (see also Supplementary Fig. 3 and Supplementary Movies 1 and 2; see Supplementary Fig. 4 for average beta band embeddings). Functionally related ROIs tended to group together, and these ROI groups segregated within embedding space. For example, auditory cortical and prefrontal ROIs were at opposite ends of dimension 1, as were visual cortical (ITGP, ITGM, LingG, FG) and prefrontal ROIs. Parietal and limbic ROIs were at opposite ends of dimension 2, and auditory and visual ROIs were maximally segregated along dimension 4. By contrast, some ROIs [e.g., STGA, anterior and middle portions of middle temporal gyrus (MTGA, MTGM), middle cingulate (CingM)] were situated in the interior of the data cloud.

One advantage of applying DME to electrophysiological data is the opportunity to examine features of the embeddings that are band-specific. DME applied to bands other than gamma produced similar embeddings. Inter-ROI distances were similar for adjacent bands (*r* ≥ 0.82), and even for non-adjacent bands (*r* ≥ 0.67; Supplementary Fig. 5). Thus, DME identified some organizational features of cortical networks that were not band-specific.

The mean correlation between inter-ROI embedding distances for original versus bootstrapped data was high in each band (r = 0.91, 0.85, 0.87, 0.88, and 0.86 for high gamma, gamma, beta, alpha, and theta, respectively). These analyses suggest that DME offers a robust approach to exploring functional geometry.

### DME elucidates fine-scale functional organization beyond anatomical proximity

The connectivity metric employed here discards components exactly in phase between two brain regions, mitigating the influence of volume conduction[68]. However, brain areas that are anatomically close to each other are often densely interconnected[69–71]. Thus, anatomical proximity is expected to contribute to the observed functional geometry. Overall, however, anatomical proximity explained only 14% of the variance in embedding distance derived from gamma band connectivity (mean adjusted *r*^2^ = 0.14 for regressions between anatomical and embedding Euclidean distance, calculated separately for each ROI). Anatomically adjacent ROIs that were separated in embedding space for gamma band included STGA and STGM, temporal pole (TP) and the rest of the anterior temporal lobe (ATL), and InsA and InsP. Similar results were obtained for embeddings derived from beta band data (Supplementary Figure 4). Thus, the embedding representation elucidates organizational features beyond anatomical proximity.

### Planum polare (PP) and posterior insula (InsP) are functionally distinct from other auditory cortical ROIs

The grouping of canonical auditory ROIs is apparent in Figure 3b and Supplementary Figure 4, as PT, HGAL, and middle and posterior portions of the superior temporal gyrus (STGM, STGP) were all close to HGPM in embedding space. One notable exception, planum polare (PP), located immediately anterior to anterolateral Heschl’s gyrus (HGAL), segregated from the rest of auditory cortical ROIs along dimension 2 in embedding space (Fig. 3b, upper panel, lower left corner; Supplementary Fig. 4, upper left panel, lower left corner). This result is consistent with PP being a higher order auditory area.

In contrast, InsP is a region that is anatomically distant from HGPM yet responds robustly to acoustic stimuli[32], suggesting that a portion of this area could be considered an auditory region[72]. For example, InsP can track relatively fast (>100 Hz) temporal modulations, similar to HGPM[32, 73], possibly due to direct inputs from the auditory thalamus. However, InsP was functionally segregated from HGPM and was situated between auditory and limbic ROIs, consistent with the broader role of InsP in polysensory exteroceptive processing and interoception[74, 75].

### Hierarchical distinction of STSU and STSL

Unlike InsP and PP, STSU was located near early auditory regions in embedding space, and for gamma band was significantly closer to auditory cortex (core and non-core ROIs; see Fig. 1) in embedding space compared to STSL (test by permutation of STSU/STSL labels, *p* < 0.0001). In beta band, the difference in distance to auditory cortex was not significant (*p* = 0.051). This distinction between STSL and STSU is consistent with differences in their response properties reported recently[23]. Particularly, responses in STSL, but not STSU, were predictive of performance in a semantic categorization task. Those results suggest that STSL would likely be closer in embedding space to regions involved in semantic processing compared to STSU. Indeed, for gamma band, STSL was significantly closer to ROIs reported to contribute to semantic processing [inferior frontal gyrus (IFG) pars operculum/triangularis/orbitalis (IFGop, IFGtri, IFGor), TP, STGA, MTGA, MTGP, anterior and posterior portions of inferior temporal gyrus (ITGA, ITGP), anterior and posterior angular gyrus (AGA, AGP), supramarginal gyrus (SMG)][76–78] compared to STSU (test by permutation of STSU/STSL labels, *p* = 0.0011). Similar results were obtained in beta band (*p* = 0.00044)

### Organization of ROIs outside auditory cortex

The data of Figure 3b and Supplementary Figure 4 also characterize the temporal and parietal ROIs outside auditory cortex that are nonetheless part of the extended auditory network, including components of the dorsal and ventral processing streams. These ‘auditory-related’ ROIs (shades of green), were distributed along a considerable extent of all four dimensions, consistent with functional heterogeneity of these regions and their involvement in integration of sensory information from multiple modalities[79].

This heterogeneity, as well as the embedding locations of PP and STSU, suggests that DME can be used to improve the brain parcellation scheme from Figure 1. For instance, MTGA in that scheme was labeled as part of the ‘Auditory-related’ group based on its location on the lateral temporal convexity and its anatomical proximity to canonical auditory cortex. The ‘Other’ group contains a large and diverse collection of ROIs whose relationship to auditory structures and speech and language processing is unclear. A more principled approach is warranted to arrange these and other ROIs into functional groups or streams based upon their physiology. One approach to developing such a data-driven parcellation scheme is to apply hierarchical clustering to the data in embedding space.

### Hierarchical clustering identifies mesoscale-level organizational features: ROI groups and processing streams

Hierarchical clustering was applied to the first four dimensions of the embedded gamma band data shown in Figure 3. The analysis illustrated a mesoscale organization of cortical ROIs (Fig. 4) that aligned with the qualitative observations discussed above. As with any clustering scheme, the number of clusters is difficult to determine based on the data alone. In the left column of Figure 4a, we illustrate two possible thresholds yielding 5 and 9 clusters, respectively. In the 5-cluster scheme, auditory cortical ROIs (excluding PP) formed an ‘Auditory’ cluster with STSU at one end of the dendrogram. At the other end, sensorimotor ROIs and ROIs typically considered part of the dorsal auditory stream formed clusters (labeled ‘Action’ and ‘Dorsal’, respectively). The remaining two large clusters were dominated by ventral temporal and limbic ROIs and by prefrontal and mesial ROIs (colored green and blue, respectively).

**Figure 4.**
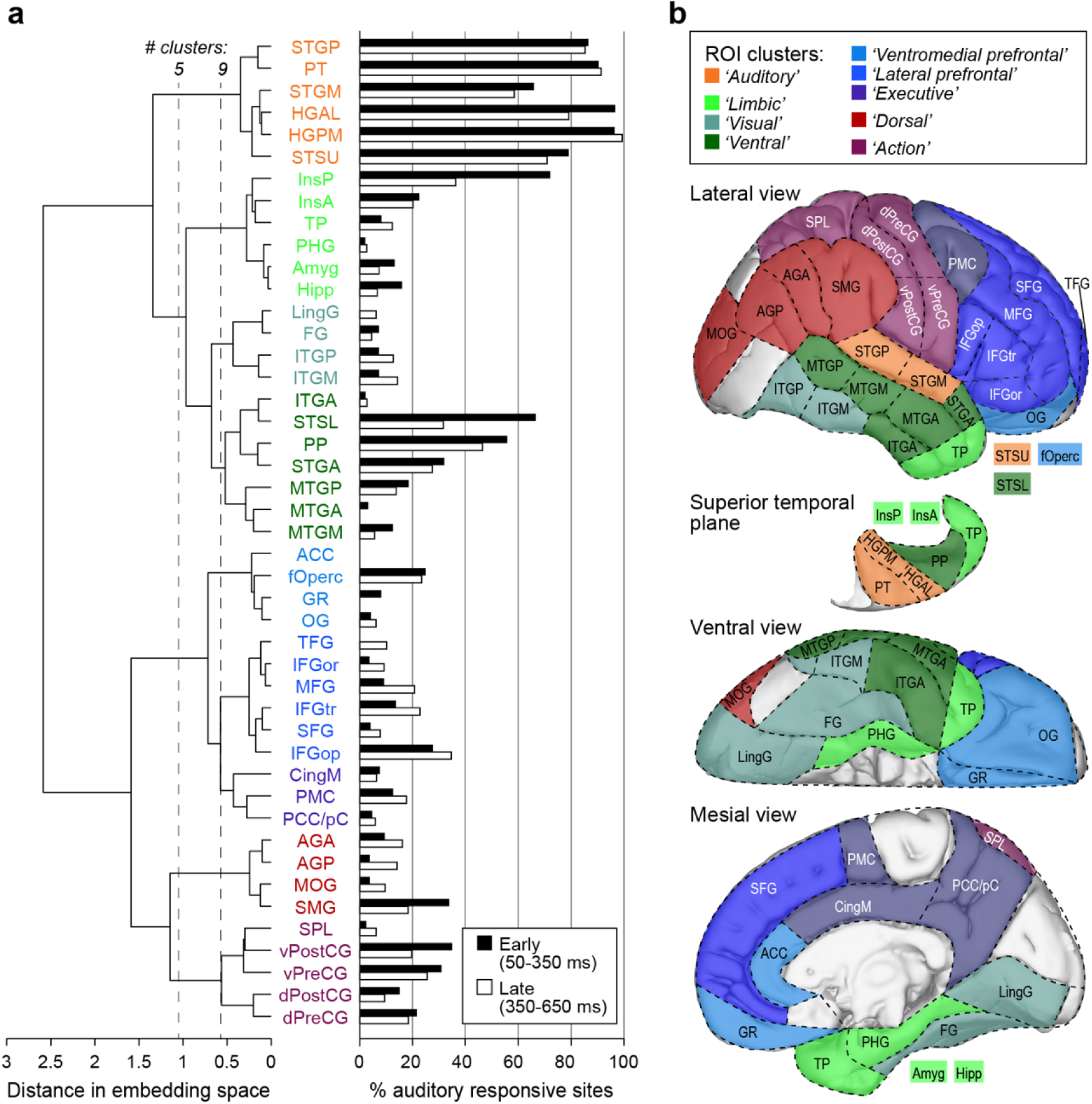
Hierarchical clustering of embedding data shown in Figure 3. **a:** Linkages between ROI groups identified using agglomerative clustering. Two thresholds are denoted (vertical dashed lines), one yielding 5 clusters and one yielding 9. ROIs are colored to indicate cluster membership. **b:** Auditory responsiveness in each ROI. Shown are percentages of sites in each ROI with early (50-350 ms after stimulus onset; black bars) and late (350-650 ms; white bars) high gamma responses to 300 ms monosyllabic words. **c:** Brain parcellation based on hierarchical clustering illustrated in **a**.

At a lower threshold, a 9-cluster scheme emerged. The ventral temporal/limbic cluster divided into three distinct clusters. One of these (‘Limbic’) included ROIs traditionally considered part of the limbic system [parahippocampal gyrus (PHG), amygdala and hippocampus], as well as TP and the insula. A second (‘Visual’) included ROIs in the ventral visual stream, and a third (‘Ventral’) consisted of ROIs typically considered part of the ventral auditory stream. Similarly, the prefrontal cluster divided into three distinct clusters (‘Ventromedial prefrontal’, ‘Lateral prefrontal’, and ‘Executive’). Thus, the hierarchical clustering analysis revealed a segregation of ROIs in embedding space that aligned with known functional differentiation of brain regions. Further, we can use this analysis to expand our understanding of hierarchical relationships among clusters. For example, the ‘Auditory’ cluster was distinct from other clusters primarily in the temporal lobe, but is closer to the ‘Limbic’ cluster than ‘Ventral’ or ‘Visual’.

Results of hierarchical agglomerative clustering applied to data from all five frequency bands are shown in Supplementary Figure 6. The color scheme for the ROIs is based on the gamma band results to provide a reference for similarity and difference across bands. Auditory cortical ROIs consistently clustered together, though the specific membership of that cluster varied slightly in alpha- and beta bands. Sensorimotor ROIs consistently clustered together, usually at a considerable distance from auditory ROIs, though in high gamma band dorsal and ventral sensorimotor ROIs were separated. Prefrontal and mesial ROIs tended to cluster together in all bands, albeit at variable overall position relative to auditory and sensorimotor ROIs. PP tended to cluster with anterior temporal lobe structures, and TP with limbic structures, regardless of frequency band. Thus, the temporal scale of neuronal signaling contributes importantly to establishing the structure of functional networks, consistent with previous results[10, 11, 80-82].

We evaluated the robustness of this clustering scheme in our dataset by calculating stability as the median normalized Fowlkes-Mallows index[83] across bootstrap iterations. The index varies between 0 (random clustering across iterations) and 1 (identical clustering across iterations). The results of the analysis as a function of threshold and frequency band indicated that stability was not strongly dependent on either band or threshold, especially for 5 or more clusters (Supplementary Fig. 7a). We also calculated cluster-wise stability as a function of the number of clusters for gamma band using the Jaccard coefficient[84]. Stability varied across threshold and clusters (Supplementary Fig. 7b). Notably, the auditory cluster was the most stable for gamma band data for both the *n_Clust_* = 5 and 9 results illustrated in Figure 4. By contrast, the ‘Executive’ cluster for *n_Clust_* = 9 was the least stable of the group.

In addition to these resting state recordings, most participants engaged in additional experiments investigating representation of acoustic stimuli in the brain[23, 85–87]. We used these data to evaluate auditory responsiveness of each recording site (Fig. 4a, right column) and compare these response profiles to the clustering results of Figure 4a (left column). As expected, ROIs in the auditory cluster exhibited consistently high responsiveness to auditory stimuli, while visual ROIs did not. By contrast, some clusters exhibited mixed responsiveness (e.g. InsP in the limbic cluster), possibly indicating ROIs that serve as nodes bridging auditory and other brain networks.

A brain parcellation scheme based on the gamma band clustering results is illustrated in Figure 4c. We note that as for other parcellation schemes based on functional connectivity (e.g., [2, 88]), the specific threshold that is most relevant and useful depends on the questions being asked and the sample size available for hypothesis testing.

### DME identifies mesoscale topological features of cortical networks

In a network, ‘global hubs’ integrate and regulate information flow by virtue of their centrality and strong connectivity; spokes send and receive information to/from these hubs. Identification of these nodes is critical for understanding the topology of brain networks [89], yet there is ongoing debate about effective methods for identifying hubs and spokes[90]. Here, we propose a novel approach to use DME to identify global hubs versus spokes. First, we note that the closer an ROI is to the center of the data cloud in embedding space, the more equal is its connectivity to the rest of the network. A simulated example is illustrated in Figure 5a, which depicts a network of five ROIs, with one serving as a global hub (Fig. 5a, left panel, green). The network structure can also be represented as an adjacency matrix, wherein the hub ROI has strong connectivity with other ROIs (Fig. 5a, middle panel). In embedding space, this ROI occupies a central location, with the other four serving as spokes, i.e., nodes that interact with each other through the central hub (Fig. 5a, right panel).

**Figure 5.**
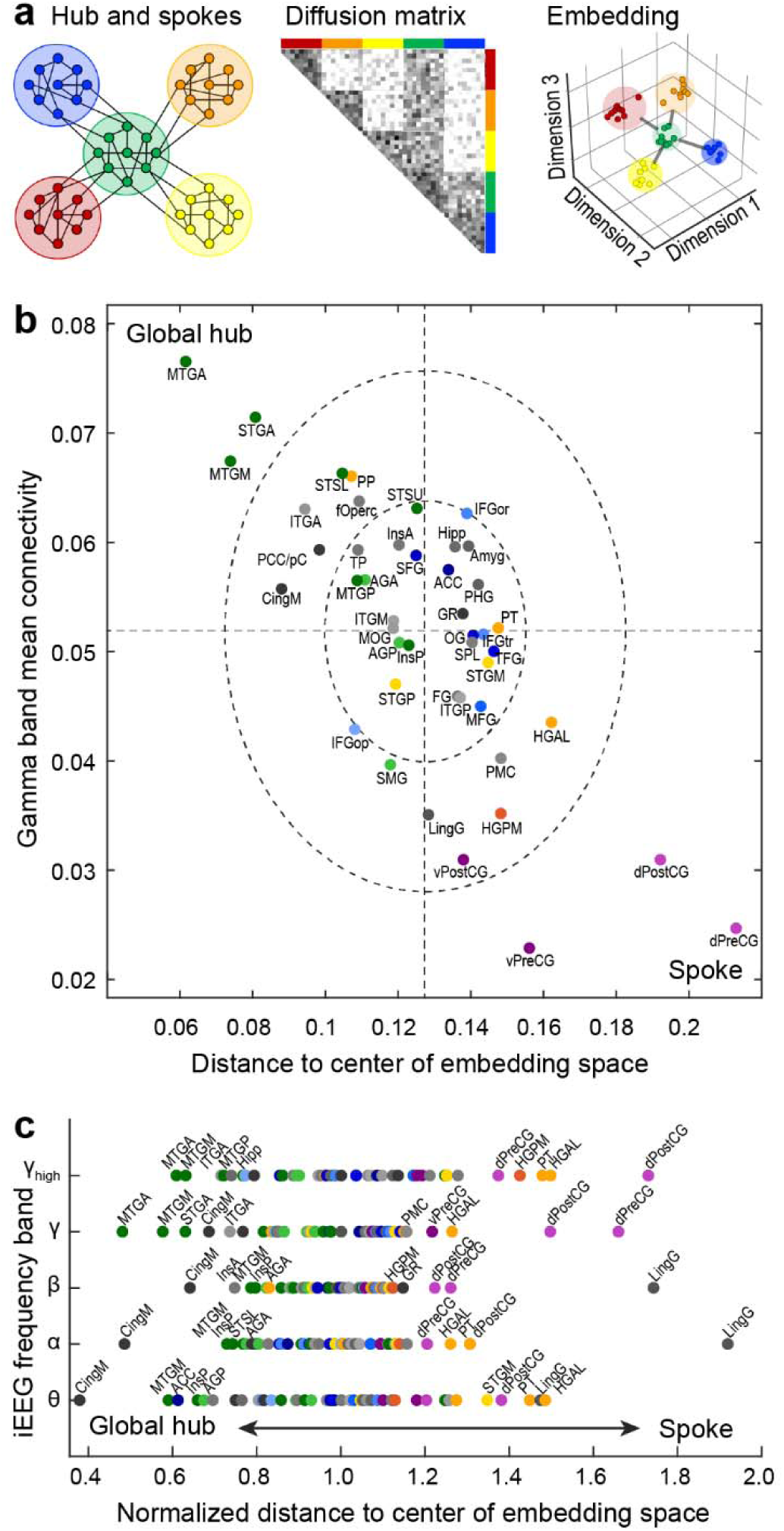
Identification of network hubs. **a:** Schematic example illustrating the central positioning of global hubs in embedding space. **b:** ROIs from average embedding are plotted according to their mean connectivity to the rest of the network versus their Euclidean distance to the centroid of the data cloud in the first four dimensions of embedding space. Dashed lines denote across-ROI means. Dashed ellipses represent 1 and 2 standard deviations from the mean. **c:** Distance to center of embedding space for each ROI in the five studied frequency bands. Distances are normalized to the median distance within each band to allow for comparison across bands.

However, a node’s proximity to the center of the data cloud reflects the *homogeneity* of its connectivity to the rest of the network, not necessarily the strength of that connectivity. In theory a node could appear at a central location if it is weakly but consistently connected to all other nodes. To determine whether this occurs in our dataset, we computed the Euclidean distance from the center of embedding space and mean connectivity for all of the ROIs in Figure 3b. We show in Figure 5b a strong inverse relationship between these two measures. ROIs close to the center of embedding space also exhibited strong mean connectivity, suggesting that global hubs can be identified in these data using distance from the center of embedding space alone.

The identity of global hubs, and the extent to which specific nodes act as global hubs, varied across frequency bands. In the high gamma and gamma bands, ROIs presenting most strongly as global hubs included MTGA, STGA, and MTGM. ITGA, CingM, posterior cingulate/precuneus (PCC/pC), PP, fOperc, and STSL also exhibited hub-like properties. By contrast, the ROIs that were farthest from the center of embedding space were mostly unimodal sensory and motor regions, consistent with their roles as spokes in the network. The positioning of these ROIs in embedding space and their roles as spokes are also indicated by the position of these ROIs at the edges of the 1-D representation depicted in the dendrograms of Figure 4 and Supplementary Figure 6.

In lower frequency bands, by contrast, CingM along with MTGM, InsP, and InsA, presented most strongly as global hubs, with the addition of ACC in the theta band. These results are consistent with network organization depending on temporal scale, and suggests that mesial cortical structures regulate information flow on slower time scales, consistent with previous reports[10]. Thus, DME can identify band-specific topological features critical to information flow within cortical networks.

### Differences between language-dominant and non-dominant hemispheres are not specific to auditory-responsive and language-specialized ROIs

On a macroscopic scale, speech and language networks are lateralized in the human brain, with nearly all right-handed and most left-handed individuals left hemisphere language-dominant[91]. However, both hemispheres are activated during speech processing[33, 39, 56, 92], and the extent to which lateralization is reflected in asymmetries in the organization of resting state auditory networks is unclear. We hypothesized that that we would observe asymmetry in gamma band data, specifically that ROIs would be in different positions in embedding space in the language-dominant versus non-dominant hemisphere. We investigated this issue by comparing the functional geometry of cortical networks derived from participants with electrode coverage in the language-dominant (*N* = 24) versus non-dominant (*N* = 22) hemisphere. ROIs in the two hemispheres exhibited a similar functional organization in embedding space (Supplementary Fig. 8). Permutation analysis indicated that for gamma band, the positions of ROIs in embedding space were not significantly different between dominant and non-dominant hemispheres (all *p*-values > 0.05). Furthermore, there was no significant correlation between the change in position in embedding space and either early or late auditory responsiveness (early: *p* = 0.94; late: *p* = 0.86; Fig. 6a). Similar results were obtained in exploratory analyses of beta band data, though one ROI (MTGP, *p* = 0.013) did survive false discovery rate (FDR) correction for difference in position between the two hemispheres.

**Figure 6.**
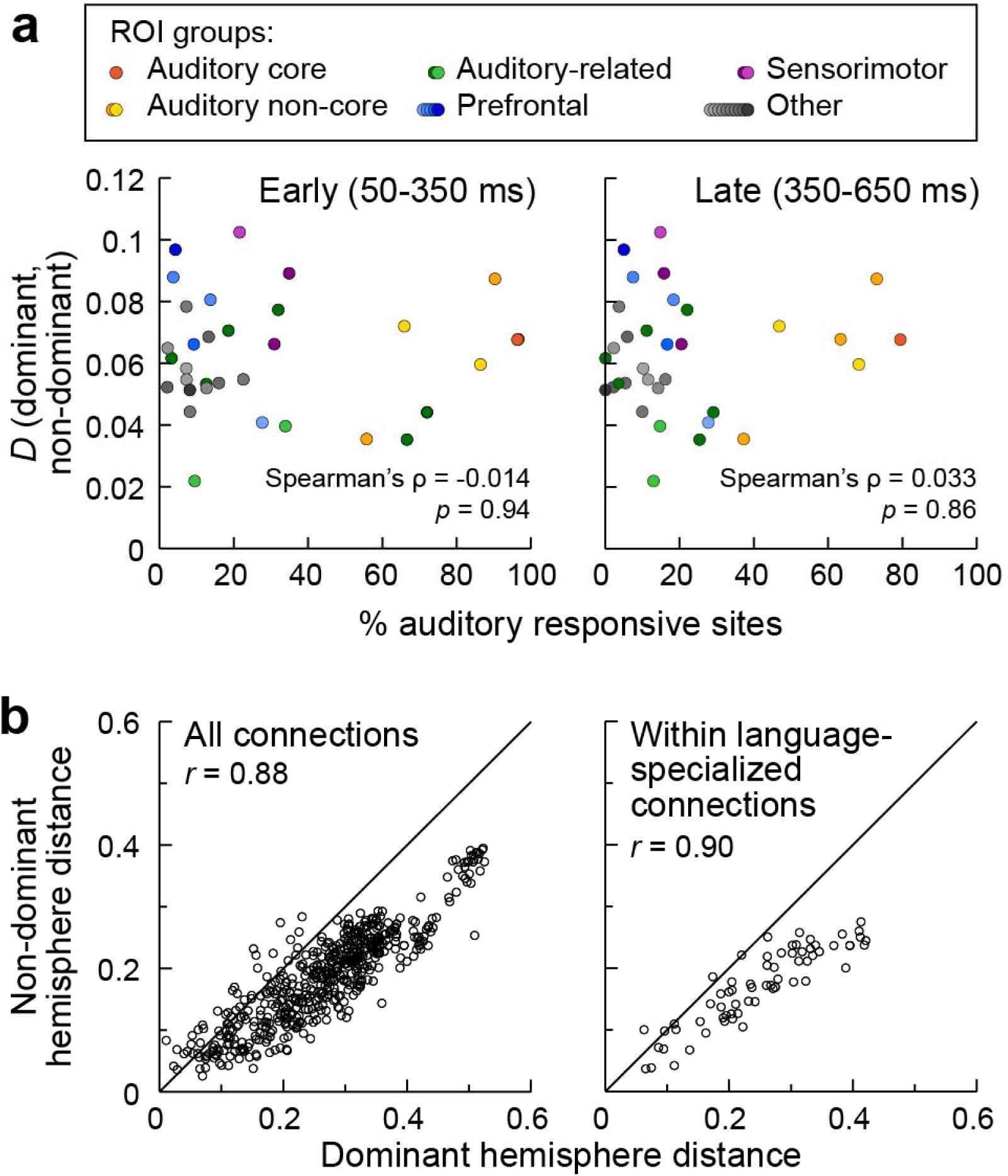
Hemispheric asymmetries in RS connectivity are not driven by auditory-responsive and language-specialized ROIs. Inter-ROI distances in embedding space for non-dominant versus dominant hemisphere participants. **a:** Comparison between the change in position in embedding space from dominant to non-dominant hemisphere and the auditory responsiveness of individual ROIs. Two-tailed Spearman’s rank tests did not reveal a significant correlation between ROI asymmetry and percentage of either early or late auditory responsive sites within the ROI (left and right panel, respectively). **b:** Pairwise distances between all ROIs and between ROIs involved in speech and language perception and production (PT, PP, STSL, STGP, STGM, STGA, SMG, AGA, PMC, PreCG, IFGop, IFGtr) are shown in the left and right panel, respectively. Note that after splitting the data into the two subsets (dominant and non-dominant) STSU did not meet the inclusion criteria for analysis presented in the right panel (see Methods, Supplementary Table 2).

We also analyzed inter-ROI distances to determine whether functional interactions between ROIs were different in the two hemispheres. For gamma band, pairwise inter-ROI distances in embedding space, calculated separately for dominant versus non-dominant hemisphere, were highly correlated (*r* = 0.88), with no obvious outliers (Fig. 6b, left panel). The data shown in Figure 6a have a slope <1, indicating that inter-ROI distances are consistently longer in the dominant hemisphere compared to the non-dominant hemisphere (*p* = 0.0032). This multiplicative scaling of the distances is consistent with the data occupying a larger volume in embedding space for the dominant versus non-dominant hemisphere, suggesting a greater functional heterogeneity for the language-dominant side of the brain. After accounting for this multiplicative scaling effect, following FDR correction, there were no specific inter-ROI distances that were significantly different between the two hemispheres. Similar results were obtained in exploratory analyses of beta band data (pairwise inter-ROI distances*r* = 0.79; longer inter-ROI distances in dominant hemisphere *p* = 0.0071; no pairwise distances significant after FDR correction).

When considering ROIs specifically involved in speech and language comprehension and production [PT, PP, STSL, STGP, STGM, STGA, SMG, AGA, premotor cortex (PMC), precentral gyrus (PreCG), IFGop, IFGtr][36, 42, 93], the correlation in pairwise inter-ROI distances in embedding space was also high (*r* = 0.90; Figure 6b). Furthermore, the data in Figure 6b exhibited a similar multiplicative scaling as observed for all the ROIs shown in Figure 6a. Indeed, the slope for the data in Figure 6b was indistinguishable from the slope for the data in Figure 6a (*p* = 0.93). Similar results were obtained in exploratory analyses of beta band data (pairwise inter-ROI distance correlations *r* = 0.76; difference between speech and language ROIs versus others *p* = 0.35). Thus, hemispheric asymmetry of functional organization specific to speech and language networks was not detectable in RS connectivity.

### Comparison to embeddings derived from RS-fMRI data

So far, we’ve presented results at multiple spatial scales based on intracranial electrophysiology. However, these intracranial recordings sample the brain non-uniformly and sparsely as dictated by clinical considerations. This feature presents problems at two spatial scales: first, cortical regions are not sampled uniformly (with some not sampled at all). Second, ROIs are not sampled uniformly across their volume. To examine the impact of these sampling issues, we compared iEEG-based DME to DME applied to RS-fMRI data available in a subset of ten participants.

We first tested the consistency of functional geometry derived from the two modalities in the same participants (Fig. 7). Connectivity matrices were constructed based on RS-fMRI data from voxels located at iEEG recording sites and grouped into the same ROIs as in Figure 1. The iEEG and fMRI embeddings averaged across participants were qualitatively similar (Fig. 7a, b), and the overall organization derived from this subset was consistent with that observed in the full iEEG dataset (cf. Fig. 3b). Inter-ROI distances in the fMRI and iEEG embedding spaces were correlated (Fig. 7c). These correlations varied across band, with highest correlations for gamma and high gamma band envelopes (*r* > 0.45; Fig. 7d, line and symbols), consistent with previous reports[80, 94].

**Figure 7.**
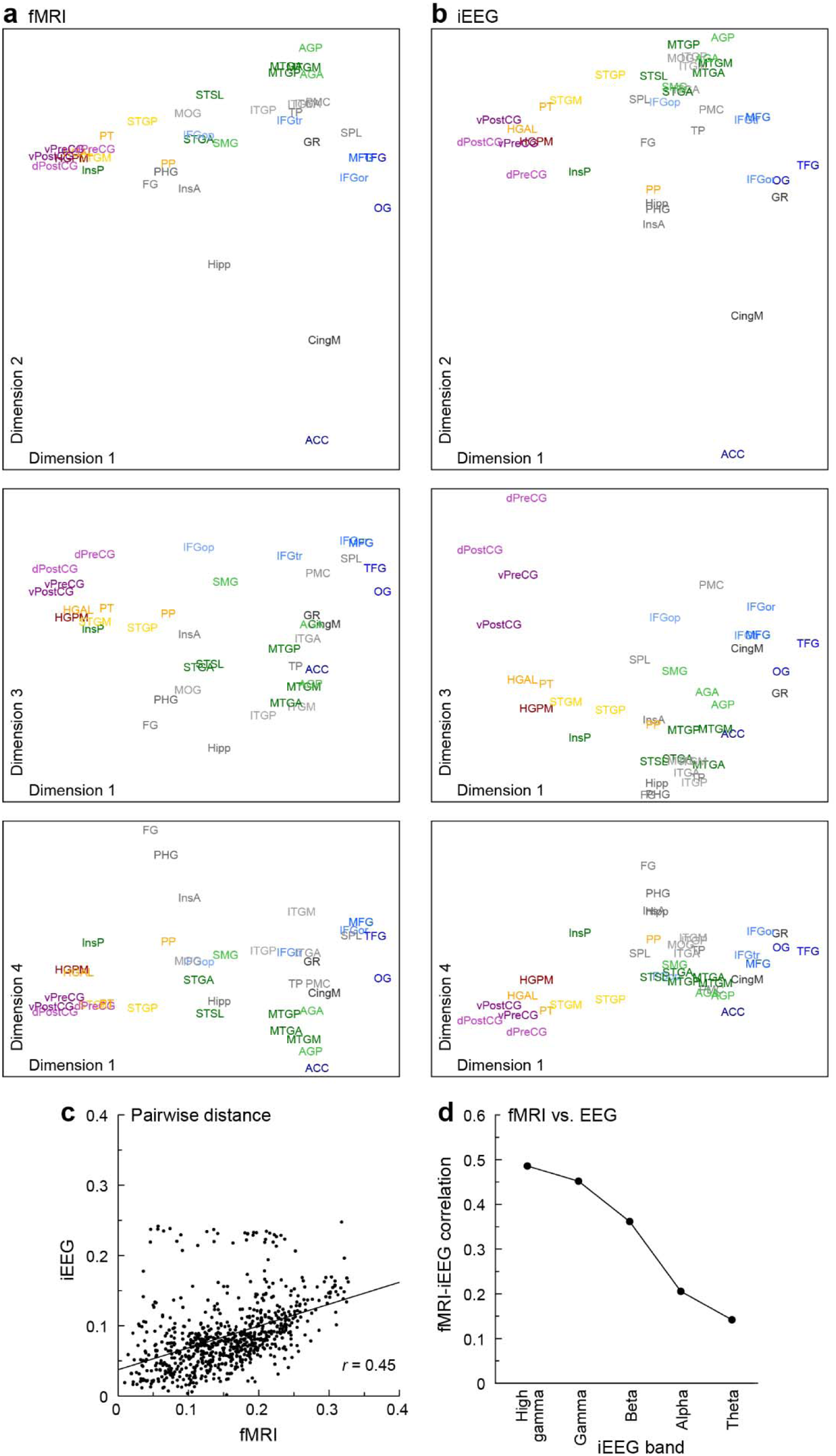
Comparison of iEEG and fMRI connectivity data in embedding space. **a:** Participant-averaged embeddings for iEEG (gamma band power envelope correlations). **b**: Participant-averaged embeddings for fMRI. **c:** Inter-ROI embedding distances computed from the data in **a** and **b**. **d:** Summary of distance correlations at each frequency band.

The analysis presented in Figure 7 provide a context for using fMRI data to address questions regarding the effects of limited, non-uniform sampling. We used a standard parcellation scheme developed for fMRI data (Schaefer-Yeo 400 ROIs;[88]) rather than the iEEG parcellation scheme introduced in Figure 1.

The first question we addressed was the effect of non-uniformly sampling only a subset of brain regions. For each participant, embeddings were derived from RS-fMRI connectivity matrices computed from all cortical ROIs (Fig. 8a, “Full fMRI”, first column). From these embeddings, we selected only points in embedding space corresponding to ROIs sampled with iEEG (Fig. 8a, “Full fMRI (iEEG subset)”, second column). We also computed embeddings for each subject from only the fMRI ROIs sampled with iEEG in that subject [“Partial fMRI (ROI level)”, Fig. 8a, 3rd column]. We compared these embeddings to the “Full fMRI (iEEG subset)” embeddings by computing the correlation between inter-ROI distances (Fig. 8b). Although the scale of the embeddings was different for the full fMRI versus partial fMRI data (because the number of dimensions was different), the two were highly correlated (median *r* = 0.90; Fig. 8c). Thus, embeddings constructed from the portion of the brain sampled by iEEG were quite similar to embeddings derived from the whole brain.

**Figure 8.**
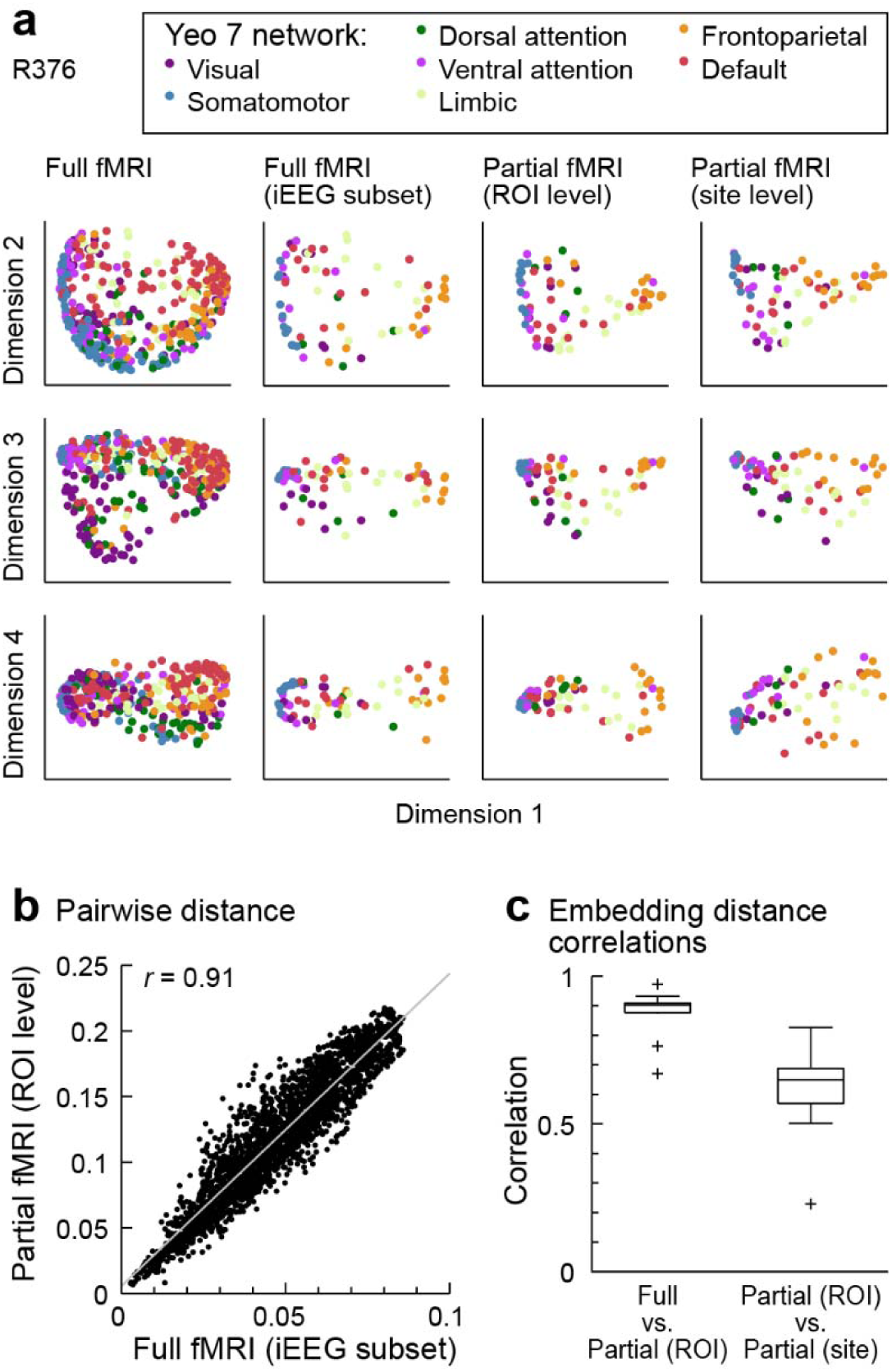
Comparison of embeddings derived from full fMRI connectivity matrices and connectivity matrices computed using only ROIs sampled with iEEG. **a:** Data in the first four dimensions of embedding space for a single participant. Shown are embeddings of all derived from the full RS-fMRI connectivity matrix (1st column); the subset of the data points in the 1st column corresponding to ROIs sampled via iEEG (2nd column); and embeddings derived from connectivity matrices including only the ROIs sampled via iEEG, calculated by averaging across the entire ROI (3rd column), and calculated based on the specific recording sites in that participant (4th column). **b:** Comparison of embedding distances calculated from the full fMRI embedding (i.e., data in **a**, 2nd column) versus distances calculated from the partial fMRI embedding (i.e., data in **a**, 3rd column). **c:** Summary across participants of distance correlations between full fMRI embeddings versus partial embeddings calculated based on the entire ROI (left: “Full vs. Partial (ROI)”) and between partial embeddings calculated based on the entire ROI versus those calculated based on recording sites [i “Partial (ROI) vs. Partial (site)”].

The second question we addressed was the effect of representing an entire ROI by sparse sampling with a limited number of electrodes. We computed embeddings from the voxel averages across entire ROIs in each participant [“Partial fMRI (ROI level)”, Fig. 8a, 3rd column] and from averages of the voxels in grey-matter spheres around iEEG recording sites [“Partial fMRI (site level)”, Fig. 8a, rightmost column]. ROI- and site-level embedding distances were strongly correlated (median *r* = 0.65; Fig. 8c).

Thus, sparse sampling within an ROI had a greater impact on estimates of functional geometry than limited sampling of the complete set of ROIs. Overall, however, ROIs were faithfully represented in embedding space even when DME was based on a small number of locations within ROIs. Taken together, these results indicate broad consistency between functional organization derived from iEEG and fMRI and the robustness of this approach to sparse sampling afforded by iEEG recordings.

## Discussion

### Organization of auditory cortical networks

We have shown that DME applied to iEEG data can be used to characterize the organization of the human auditory cortical hierarchy at multiple spatial scales. We demonstrate methodology for testing specific hypotheses about gamma band data at each of these scales using DME. We also use exploratory analyses (e.g., hierarchical clustering, analyses of other frequency bands) to generate data-driven hypotheses for study using future data sets.

### Investigating cortical network organization using resting state data

The results presented here are based on analysis of resting state (i.e., task-free) data. Relationships between brain signals recorded at different locations derive from synaptic connections between neurons in those locations. Thus, these data provide valuable information about the underlying brain organization despite the absence of a task or a controlled sensory stimulus. The same areas that are co-activated during sensory processing exhibit resting state connectivity with each other, and resting state networks map onto relevant behavioral and task-related domains [2, 3]. Numerous previous studies based on BOLD fMRI have used analyses of resting state activity to gain insight into the organization of human brain networks and how this organization is altered due to brain disorders, during development and ageing, and in response to pharmacological treatments [22, 26, 95–99].

A key advantage of resting state analyses is that they are based data that is far more stationary compared to data derived from task-based experiments. In the case of sensory regions, this avoids a confound inherent to investigations of connectivity in the presence of a stimulus, which itself would produce correlated activity at directly driven sites. That is, two unconnected sites in the brain driven by the same sensory stimulus will exhibit apparent connectivity solely due to the stimulus, in the absence of a physical connection between the sites. Additionally, resting state analyses typically can draw on considerably more data than is available from task-based experiments, allowing for better estimates of connectivity.

As in previous studies, we provide a snapshot of the organization of these networks, corresponding to a static representation. However, these networks are dynamic due to short- and long-term plasticity driven both by internal (e.g., changes in arousal state) and external factors (e.g., sensory stimulation and directed behavior). The organization derived from studies such as this one provides a framework for understanding these dynamics.

### Frequency band-specific properties of cortical networks

Previous reports have shown that cortical networks defined by functional or effective connectivity derived from electrophysiological data exhibit organizational structure that depends on the frequency band being analyzed. This manifests in two ways relevant to the results presented here. First, canonical resting state networks originally identified using resting state BOLD fMRI data vary across band in the strength of within-network connectivity and in the relationship between electrophysiological- and fMRI-derived connectivity networks [10, 80]. Second, detailed analyses of the relationship between anatomical projection patterns and functional or effective connectivity indicate that especially in auditory and visual cortical hierarchies, feedforward information streams rely on connectivity primarily in gamma and high-gamma bands, while lower frequency bands (alpha, beta) underlie feedback connectivity[11–19]. Based on these previous reports, we suggest that the gamma band organization that is the focus of the current report reflects feedforward connectivity. Indeed, the auditory responsiveness profile depicted in Figure 4b is most strongly predicted by clustering analysis applied to gamma and high-gamma band data in embedding space. Results for other bands differed from gamma band results especially in the identity of network hubs (Figure 5), where middle cingulate cortex emerged as the ROI with the most pronounced ‘hubness’. By contrast, the overall organization features considered at various spatial scales did not differ strongly between bands, suggesting that while temporal scale is an important contributor to network organization, functional connectivity on these different scales tends to overlap. In the case of comparisons between feedforward and feedback networks, this is consistent with the tendency of cortical areas to be coupled bidirectionally[100].

### Fine scale: Organization of auditory cortex

At a fine spatial scale, previous work in non-human primates has defined over a dozen auditory cortical fields based on cytoarchitectonics, connectivity, and response properties[101]. By contrast, there is no consensus on how auditory cortex is organized in humans, with multiple parcellations based on cytoarchitectonics, tonotopy or myeloarchitecture[102–105]. Our results contribute to this body of knowledge by showing that several superior temporal ROIs including core auditory cortex (HGPM) and putative auditory belt and parabelt areas (PT, HGAL, STGM)[102, 105] group together in embedding space across all frequency bands. Thus, in spite of their diversity in processing of specific features of acoustic signals, these ROIs are positioned at a similar level in the auditory cortical hierarchy. Other regions, such as STGP and STSU, group with these cortical ROIs in theta, gamma, and high-gamma, but not in alpha and beta. For gamma band, proximity of STGP and STGM to HGPM in embedding space is consistent with previous studies that interpret these regions as relatively early non-core auditory cortex[29, 106, 107]. By contrast, although PP is anatomically close and connected to HGPM[108], for both gamma and beta band it was not close to HGPM in embedding space. PP is distinguished among auditory cortical regions for its syntactic-level language processing[30] and its preferential activation by music, which has a strong affective component[31]. This functional differentiation is reflected in its segregation from the group of auditory cortical ROIs in embedding space.

### Fine scale: Functional differentiation between STSU and STSL

The superior temporal sulcus is a critical node in speech and language networks linking canonical auditory cortex with higher order temporal, parietal, and frontal areas[22, 33–37]. Previous studies have shown that STSU and STSL differ in cytoarchitecture[109] and have distinct responses to speech[27, 57, 110, 111]. A recent iEEG study demonstrated enhanced, shorter-latency responses to speech syllables in STSU compared to STSL[23]. STSU is traditionally not considered part of canonical auditory cortex (but see[103]), yet it was located close to auditory cortical ROIs in embedding space in gamma band. STSL, by contrast, was closer in embedding space to semantic ROIs in both beta and gamma band. This is consistent with iEEG evidence that responses in STSL, but not STSU, correlated with performance on a semantic categorization task[23]. The regions specifically involved in semantic processing is a current topic of debate, with multiple competing models[21, 76–78]. We defined a list of semantic processing regions by combining across these models. Taken together, the results firmly place STSU and STSL at different levels of the auditory cortical hierarchy defined by gamma band connectivity.

### Mesoscale: Functional and theoretical framework of a limbic auditory pathway

Multiple lines of evidence support a pathway linking auditory cortical and limbic structures[112–115] that subserves auditory memory[45, 48, 49] and affective sound processing[116]. The data presented here contribute to our understanding of this pathway. Clustering analysis identified a set of ROIs including structures classically labeled as limbic (PHG, Amy, Hipp) as well as insula (InsP, InsA) and TP positioned close to the auditory cluster in embedding space for both gamma and beta bands (Fig. 4; Supplementary Fig. 4). This suggests a close functional relationship that could form the basis for a limbic stream. InsP, with strong auditory responsiveness and overlapping response properties with HGPM, is likely involved in the transformation of auditory information in auditory cortex to affective representations in InsA[32]. Thus, InsP could serve as critical linking node between auditory and limbic structures.

TP is involved in semantic processing[21, 30] and auditory memory[117], in particular the representation and retrieval of memories for people, social language, and behaviors (‘social knowledge’)[118]. Tight clustering of TP with limbic ROIs in embedding space is consistent with its previously reported functional association with limbic cortex[119, 120], with which TP shares key features of laminar cytoarchitecture and strong connectivity[121]. We suggest that the organization depicted in Figures 3 and 4, combined with evidence for bidirectional information sharing between auditory cortex and limbic areas, merits the identification of a third auditory processing stream alongside the dorsal and ventral streams[38, 122]. This ‘limbic stream’ would underlie auditory contributions to affective and episodic memory processing.

### Mesoscale: Ventral and dorsal streams linking auditory and frontal cortex

Current models of speech and language processing posit the existence of ventral and dorsal processing streams linking non-core auditory cortex with PMC and inferior frontal gyrus via several distinct anatomical pathways encompassing temporal, parietal, and frontal cortex[36, 38–40]. Despite substantial experimental evidence supporting these models, there is a lack of consensus on the specific functions subserved by the two streams. For example, while there is consensus that the ventral stream subserves auditory object identification (“what” processing), the dorsal stream has been envisioned to subserve spatial processing (“where”[38]) and audiomotor processing[39]. There is a parallel debate about the specific cortical regions comprising the two streams.

As broadly predicted by these models, temporal and parietal ROIs segregated in embedding space in the analysis presented here (Fig. 3b, 4; Supplementary Figs. 4 & 6). Across frequency bands, we observed a cluster that included STSL and ATL ROIs, in conformity with the ventral auditory stream proposed by Hickok and Poeppel[39] and Friederici[40]. By contrast, the cluster that included SMG, AGP, and AGA aligned with the dorsal processing stream as proposed by Rauschecker and Scott[38]. The proximity of these dorsal ROIs to sensorimotor ROIs is consistent with sensorimotor contributions to dorsal stream processing[43, 123]. Association of FG and MOG with the ventral and dorsal clusters, respectively, likely represents the sharing of information across sensory modalities. For example, visual information has been shown to contribute to processing in the ventral (“what”) pathway[124, 125].

A previous fMRI-based DME study found that primary sensory and default mode ROIs segregated along the first dimension in embedding space[25]. Coverage of mesial cortex in our dataset was limited, precluding a direct comparison. However, the striking separation between auditory and prefrontal cortex in embedding space shown here indicate that the current results align well with the previous report. This separation places auditory and frontal regions at opposite ends of a cortical hierarchy, linked by ventral and dorsal processing streams[38–40].

### Mesoscale: Network hubs

Hubs in brain networks play a critical role in integrating distributed neural activity[89, 126]. In the present analysis, global hubs were characterized by their central location within embedding space (Fig. 5). In the gamma band, these hubs included STGA and MTGA, both components of the ATL. Previous reports indicate that ATL serves as a transmodal hub, transforming sensory domain-specific to domain-general representations[21, 127, 128] and playing a central role in semantic processing and social memory[21, 118, 129]. MTGM also appears as a global hub, even though it is anatomically distinct from the ATL. Interestingly, patients with semantic dementia have ATL degeneration[130, 131], but the damage is often more widespread and can include MTGM[132].

Cingulate cortical ROIs (CingM, ACC) and insula were identified as hubs in lower frequency bands. CingM and ACC are described as transmodal and are active during a wide array of emotional and cognitive processes[133, 134], both consistent with their previous characterization as network hubs[126]. The identification of hubs specific to each frequency band supports the model in which the temporal scale of communication in the brain supports distinct functional networks[80–82, 135]. Also consistent with this model is the frequency band-specific correspondence between iEEG and fMRI connectivity observed here (Fig. 7d) and in previous reports[80, 94]. Strong correspondence between BOLD fMRI connectivity and higher frequency band envelope correlations in iEEG are observed, while the correspondence for theta and alpha bands is usually still positive but lower in magnitude. Of note, this frequency dependence of connectivity is distinct from previous observations of a frequency-dependent correspondence between iEEG power and BOLD fMRI signal magnitude[136]. This relationship for connectivity also depends on brain location[80]. Because we did analyze the relationship between fMRI and iEEG in a region-specific manner, the results presented here represent an average analysis over all sampled brain regions.

Unlike other ATL structures, TP does not present as a global hub in any frequency band (Fig. 5c). The close association of TP with limbic structures in embedding space suggests that TP mediates interactions between transmodal integration centers in the ATL and structures subserving memory functions. More broadly, the heterogeneity of ATL ROIs in terms of their global hub-like connectivity profiles conforms to the observation that the terminal fields of white matter tracts converging in the ATL only partially overlap[21, 137, 138].

### Macroscale: Hemispheric lateralization

Although speech and language networks are classically described as highly lateralized, imaging studies have demonstrated widespread bilateral activation during speech and language tasks[52–54]. Indeed, a recent fMRI study showed RS connectivity patterns in lateral temporal cortex that were comparable between left and right hemispheres[6]. We found evidence for hemispheric differences in RS cortical functional organization based on analysis of all sampled brain regions, with inter-ROI distances being systematically greater in embedding space for the language-dominant hemisphere (Fig. 6b). This is consistent with greater inter-regional heterogeneity in that hemisphere compared to the non-dominant side. Importantly, the observed asymmetry could not be attributed specifically to ROIs involved in speech and language processing (Fig. 6b), nor was the difference in position in embedding space related to auditory responsiveness (Fig. 6a).

Recent studies that identified hemispheric differences in RS connectivity for the STS[22] and semantic networks more broadly[139] may reflect the general asymmetry observed here. This asymmetry may relate as well to the dichotomy between domain-specific (e.g., sensory processing) and domain-general (e.g., attention, working memory) cortical systems. In particular, studies have emphasized that domain-general systems also exhibit hemispheric laterality[140, 141], suggesting that the asymmetry observed here may reflect this broader organization feature. This does not exclude the possibility of asymmetries specific to auditory regions emerging during sensory tasks, for example reflecting hemispheric biases in spectral and temporal processing[39, 42].

### Caveats & limitations

A key concern regarding all human iEEG studies is that participants may not be representative of a healthy population. In the present study, results were consistent across participants despite differences in seizure disorder histories, medications, and seizure foci, and aligned with results obtained previously in healthy participants[25]. Another caveat is that our dataset, however extensive, did not sample the entire brain, and it was not possible to infer connectivity with unsampled regions. To address this, we applied DME analysis to fMRI data to establish that the organization of ROIs in embedding space was robust to the exclusion of unsampled ROIs. Although there was a greater effect of sparse, non-uniform sampling within an ROI, there was still considerable similarity in functional organization to embeddings derived from averages across the entire ROI

While subcortical structures (e.g., thalamus) that link sensory and higher order networks[142] were not sampled, the functional organization presented here was likely influenced indirectly by thalamo-cortical pathways[29, 143]. Previous fMRI studies of RS networks focused exclusively on cortical ROIs and did not consider the role of the thalamus and other subcortical structures. Despite this limitation, these studies have yielded valuable insights into the functional organization of the human cortical networks[1, 144].

### Concluding remarks and future directions

This study extends the DME approach to characterize functional relationships between cortical regions investigated using iEEG recordings. These data help resolve several outstanding issues regarding the functional organization of human auditory cortical networks and stress the importance of a limbic pathway complementary to the dorsal and ventral streams. These results lay the foundation for future work investigating network organization during active speech and language processing. The superior time resolution of electrophysiological data allows for dynamic connectivity analysis on time scales relevant to this processing. An important next step for this work is to adapt this analysis to scalp EEG recordings, which offer considerable advantages over fMRI in terms of accessibility and cost. While the current work focused on auditory cortical networks, this approach can be readily generalized to advance our understanding of changes in brain organization during sleep and anesthesia, disorders of consciousness, as well as reorganization of cortical functional geometry secondary to lesions.

## Materials and Methods

### Ethics Statement

Research protocols aligned with best practices recently aggregated in[145] and were approved by the University of Iowa Institutional Review Board and the National Institutes of Health; written informed consent was obtained from all participants. Research participation did not interfere with acquisition of clinically necessary data, and participants could rescind consent for research without interrupting their clinical management.

### Participants

The study was carried out in 49 neurosurgical patients (22 females) diagnosed with medically refractory epilepsy. The patients were undergoing chronic invasive electrophysiological monitoring to identify seizure foci prior to resection surgery (Supplementary Table 1). All participants except two were native English speakers. The participants were predominantly right-handed (42 out of 49); six participants were left-handed, and one had bilateral handedness. The majority of participants (35 out of 49) were left language-dominant, as determined by Wada test. Two participants were right hemisphere-dominant, and one had bilateral language dominance. The remaining 11 participants were not evaluated for language dominance; 9 of them were right-handed and thus were assumed left language-dominant for the purposes of the analysis of lateralization (see below). The participant with bilateral dominance, and the remaining two participants who did not undergo Wada test and who were left-handed were not included in the analysis of hemispheric asymmetry in Figure 6. All participants underwent audiological and neuropsychological assessment prior to electrode implantation, and none had auditory or cognitive deficits that would impact the results of this study. The participants were tapered off their antiepileptic drugs during chronic monitoring when RS data were collected.

### Experimental procedures

#### Pre-implantation neuroimaging

All participants underwent whole-brain high-resolution T1-weighted structural MRI scans before electrode implantation. In a subset of ten participants (Supplementary Table 2), RS-fMRI data were used for estimates of functional connectivity. The scanner was a 3T GE Discovery MR750W with a 32-channel head coil. The pre-electrode implantation anatomical T1 scan (3D FSPGR BRAVO sequence) was obtained with the following parameters: FOV = 25.6 cm, flip angle = 12 deg., TR = 8.50 ms, TE = 3.29 ms, inversion time = 450 ms, voxel size = 1.0 × 1.0 × 0.8 mm. For RS-fMRI, 5 blocks of 5-minute gradient-echo EPI runs (650 volumes) were collected with the following parameters: FOV = 22.0 cm, TR = 2260 ms, TE = 30 ms, flip angle = 80 deg., voxel size = 3.45 × 3.45 × 4.0 mm. In some cases, fewer RS acquisition sequences were used in the final analysis due to movement artifact or because the full scanning session was not completed. For each participant, RS-fMRI runs were acquired in the same session but non-contiguously (dispersed within an imaging session to avoid habituation). Participants were asked to keep their eyes open, and a fixation cross was presented through a projector.

#### iEEG recordings

iEEG recordings were obtained using either subdural and depth electrodes, or depth electrodes alone, based on clinical indications. Electrode arrays were manufactured by Ad-Tech Medical (Racine, WI). Subdural arrays, implanted in 36 participants out of 46, consisted of platinum-iridium discs (2.3 mm diameter, 5-10 mm inter-electrode distance), embedded in a silicon membrane. Stereotactically implanted depth arrays included between 4 and 12 cylindrical contacts along the electrode shaft, with 5-10 mm inter-electrode distance. A subgaleal electrode, placed over the cranial vertex near midline, was used as a reference in all participants. All electrodes were placed solely on the basis of clinical requirements, as determined by the team of epileptologists and neurosurgeons[146].

No-task RS data were recorded in the dedicated, electrically shielded suite in The University of Iowa Clinical Research Unit while the participants lay in the hospital bed. RS data were collected 6.4 +/- 3.5 days (mean ± standard deviation; range 1.5 – 20.9) after electrode implantation surgery. In the first 15 participants (L275 through L362), data were recorded using a TDT RZ2 real-time processor (Tucker-Davis Technologies, Alachua, FL). In the remaining 34 participants (R369 through L585), data acquisition was performed using a Neuralynx Atlas System (Neuralynx Inc., Bozeman, MT). Recorded data were amplified, filtered (0.7–800 Hz bandpass, 5 dB/octave rolloff for TDT-recorded data; 0.1–500 Hz bandpass, 12 dB/octave rolloff for Neuralynx-recorded data) and digitized at a sampling rate of 2034.5 Hz (TDT) or 2000 Hz (Neuralynx). In all but two participants, recording durations were between 10 and 18 minutes, the median was 10; in one participant duration was 6 min., and in one participant the duration was 81 min.

### Data analysis

#### Anatomical reconstruction and ROI parcellation

Localization of recording sites and their assignment to ROIs relied on post-implantation T1-weighted anatomical MRI and post-implantation computed tomography (CT). All images were initially aligned with pre-operative T1 scans using linear coregistration implemented in FSL (FLIRT)[147]. Electrodes were identified in the post-implantation MRI as magnetic susceptibility artifacts and in the CT as metallic hyperdensities. Electrode locations were further refined within the space of the pre-operative MRI using three-dimensional non-linear thin-plate spline warping[148], which corrected for post-operative brain shift and distortion. The warping was constrained with 50-100 control points, manually selected throughout the brain, which were visually aligned to landmarks in the pre- and post-implantation MRI.

Electrode locations were mapped into a common anatomical template space using a combination of surface-based and volumetric coregistration. Automated identification and parcellation of the cortical surface within T1-weighted images was carried out with FreeSurfer [149, 150]. Electrodes were assigned anatomical labels within the parcellation scheme of Destrieux et al. [151, 152], according to the label of the nearest vertex (within the T1 image space) of the cortical surface mesh generated by FreeSurfer. Labeling was visually inspected and corrected whenever the automated parcellation did not conform to expected gyral boundaries. Volumetric mapping of T1 images to the MNI-152 space relied on automated linear coregistration implemented in the fsl_anat pipeline of the FSL toolbox [153]. Electrode coordinates in MNI-152 space were obtained by applying the resulting transformation to the coordinates from the T1 image space.

To pool data across participants, the dimensionality of connectivity matrices was reduced by assigning electrodes to one of 58 ROIs organized into 6 ROI groups (see Fig. 1; Supplementary Table 2, 3) based upon anatomical reconstructions of electrode locations in each participant. For subdural arrays, ROI assignment was informed by automated parcellation of cortical gyri[151, 152] as implemented in the FreeSurfer software package. For depth arrays, it was informed by MRI sections along sagittal, coronal, and axial planes. For recording sites in Heschl’s gyrus, delineation of the border between core auditory cortex and adjacent non-core areas (HGPM and HGAL, respectively) was performed in each participant using physiological criteria[154, 155]. Specifically, recording sites were assigned to HGPM if they exhibited phase-locked (frequency-following) responses to 100 Hz click trains and if the averaged evoked potentials to these stimuli featured short-latency (<20 ms) peaks. Such response features are characteristic for HGPM and are not present within HGAL[154]. Additionally, correlation coefficients between average evoked potential waveforms recorded from adjacent sites were examined to identify discontinuities in response profiles along Heschl’s gyrus that could be interpreted as reflecting a transition from HGPM to HGAL. Superior temporal gyrus was subdivided into posterior and middle non-core auditory cortex ROIs (STGP and STGM), and auditory-related anterior ROI (STGA) using the transverse temporal sulcus and ascending ramus of the Sylvian fissure as macroanatomical boundaries. The insula was subdivided into posterior and anterior ROIs, with the former considered within the auditory-related ROI group[32]. Middle and inferior temporal gyrus were each divided into posterior, middle, and anterior ROIs by diving the gyrus into three approximately equal-length thirds. Angular gyrus was divided into posterior and anterior ROIs using the angular sulcus as a macroanatomical boundary. Anterior cingulate cortex was identified by automatic parcellation in FreeSurfer and was considered as part of the prefrontal ROI group, separately from the rest of the cingulate gyrus. Postcentral and precentral gyri were each divided into ventral and dorsal portions using the *z*_MNI_ coordinate (see below) of 40 mm as a boundary. Recording sites identified as seizure foci or characterized by excessive noise, or outside brain, were excluded from analyses and are not listed in Supplementary Table 2. Depth electrode contacts localized to the white matter were also excluded. Location within cortical white matter was determined based on visual inspection of anatomical reconstruction data (MRI sections along sagittal, coronal, and axial planes) as done in our previous studies (e.g., [62]). Electrode coverage was largely restricted to a single hemisphere in individual participants, and contacts on the contralateral hemisphere were excluded from analysis (and are not listed in Supplementary Table 2) such that all connections represent intra-hemisphere functional connectivity.

#### Preprocessing of fMRI data

Standard preprocessing was applied to the RS-fMRI data acquired in the pre-implantation scan using FSL’s FEAT pipeline, including spatial alignment and nuisance regression. White matter, cerebrospinal fluid and global ROIs were created using deep white matter, lateral ventricles and a whole brain mask, respectively. Regression was performed using the time series of these three nuisance ROIs as well as 6 motion parameters (3 rotations and 3 translations) and their derivatives, detrended with second order polynomials. Temporal bandpass filtering was 0.008–0.08 Hz. Spatial smoothing was applied with a Gaussian kernel (6 mm full-width at half maximum). The first two images from each run were discarded. Frame censoring was applied when the Euclidean norm of derivatives of motion parameters exceeded 0.5 mm[156]. All runs were processed in native EPI space, then the residual data were transformed to MNI152 and concatenated.

#### Preprocessing of iEEG data

Analysis of iEEG data was performed using custom software written in MATLAB Version 2020a programming environment (MathWorks, Natick, MA, USA). After initial rejection of recording sites identified as seizure foci, several automated steps were taken to exclude recording channels and time intervals contaminated by noise. First, channels were excluded if average power in any frequency band [broadband, delta (1-4 Hz), theta (4-8 Hz), alpha (8-13Hz), beta (13-30 Hz), gamma (30-50 Hz), or high gamma (70-110 Hz); see below] exceeded 3.5 standard deviations of the average power across all channels for that participant. Next, transient artifacts were detected by identifying voltage deflections exceeding 10 standard deviations on a given channel. A time window was identified extending before and after the detected artifact until the voltage returned to the zero-mean baseline plus an additional 100 ms buffer before and after. High-frequency artifacts were also removed by masking segments of data with high gamma power exceeding 5 standard deviations of the mean across all segments. Only time bins free of these artifact masks were considered in subsequent analyses. Artifact rejection was applied across all channels simultaneously so that all connectivity measures were derived from the same time windows. Occasionally, particular channels survived the initial average power criteria yet had frequent artifacts that led to loss of data across all the other channels. There is a tradeoff in rejecting artifacts (losing time across all channels) and rejecting channels (losing all data for that channel). If artifacts occur on many channels, there is little benefit to excluding any one channel. However, if frequent artifacts occur on one or simultaneously on up to a few channels, omitting these can save more data from other channels than those channels contribute at all other times. We chose to optimize the total data retained, channels × time windows, and omitted some channels when necessary.

On occasion, noise from in-room clinical equipment and muscle artifacts appeared in the data as shared signals across channels. These types of noise are typically broadband, and can be detected via analysis of frequencies higher than those of interest here. To remove these signals, data from retained channels were high-pass filtered above 200 Hz, and a spatial filter was derived from the singular value decomposition omitting the first singular vector. This spatial filter was then applied to the broadband signal to remove this common signal.

#### Connectivity analysis

For RS-fMRI data, BOLD signals were averaged across voxel groupings and functional connectivity was calculated as Pearson correlation coefficients. Voxel groupings were either based on the Schaefer-Yeo 400 parcellation scheme[88] in MNI-152 space, or were based on iEEG electrode location in participant space (see Fig. 1). For the latter, fMRI voxels were chosen to represent comparable regions of the brain recorded by iEEG electrodes. For each electrode, the anatomical coordinates of the recording site were mapped to the closest valid MRI voxel, *E*, and a sphere of 25 voxels (25 mm^3^) centered on *E* used as the corresponding recording site. This process was repeated for all *N* electrodes in the same ROI, and a single time series computed as the average of the fMRI BOLD signal in these *N*×25 voxels. These averages were used to compute an ROI-by-ROI connectivity matrix for RS-fMRI data. For comparisons between iEEG and fMRI embeddings, voxels were processed in participant space and ROI labels from the parcellation scheme illustrated in Figure 1 and Supplementary Table 2 were applied to the fMRI data. For comparisons between fMRI embeddings derived from all cortical ROIs versus fMRI embeddings derived from just ROIs sampled in the iEEG experiments, electrode locations were transformed from participant space to MNI-152 space, then assigned to ROIs within the Schaefer-Yeo 400 scheme.

Connectivity was measured for iEEG data using orthogonalized band power envelope correlation[68]. This measure avoids artifacts due to volume conduction by discounting connectivity near zero phase lag. Data were divided into 60-second segments, pairwise connectivity estimated in each segment, and then connectivity estimates averaged across all segments for that subject. Power envelope correlations were calculated using a method similar to [68], except time-frequency decomposition was performed using the demodulated band transform[157] rather than wavelets. This measure avoids artifacts due to volume conduction by discounting connectivity near zero phase lag. For each frequency band (theta, alpha, beta, gamma; high gamma), the power at each time bin was calculated as the average (across frequencies) log of the squared amplitude. For each pair of signals *X* and *Y*, one was orthogonalized to the other by taking the magnitude of the imaginary component of the product of one signal with the normalized complex conjugate of the other:

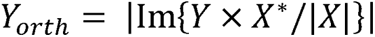

Both signals were band-pass filtered (0.2 – 1 Hz), and the Pearson correlation calculated between signals. The process was repeated by orthogonalizing in the other direction and the overall envelope correlation for a pair of recording sites was the average of the two Pearson correlations. Lastly, correlations were averaged across segments.

Connectivity matrices were thresholded prior to diffusion map embedding to reduce the contribution of spurious connections to the analysis. We balanced our desire to minimize noisy connections while maintaining a connected graph (i.e., that that there are no isolated nodes; required by DME[59]) by saving at least the top third (rounded up) connections for every row, as well as their corresponding columns (to preserve symmetry). To ensure that the graph was connected after thresholding, we also included any connections making up the minimum spanning tree of the graph represented by the elementwise reciprocal of the connectivity matrix to ensure the graph is connected.

#### ROI-based connectivity analysis

Connectivity between ROIs was computed as the average envelope correlation between all pairs of recording sites in the two ROIs.For analyses in which connectivity was summarized across participants (Fig. 3-8), we used only a subset of ROIs such that every possible pair of included ROIs was represented in at least two participants (Supplementary Table 2). This list of ROIs was obtained by iteratively removing ROIs with the worst cross-coverage with other ROIs until every ROI remaining had sufficient coverage with all remaining ROIs.

#### Diffusion map embedding

See the Appendix I for details about DME.

In brief, the functional connectivity is transformed by applying cosine similarity[25] to yield the similarity matrix **K =** [*k*(*i*,*j*)]. This matrix then normalized by degree to yield a matrix **P = D**^-1^**K**, where **D** is the degree matrix, i.e. the diagonal elements of 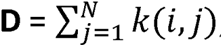, where *N* is the number of recording sites, and the off-diagonal elements of **D** are zero. If the recording sites are conceptualized as nodes on a graph with edges defined by **K**, then **P** can be understood as the transition probability matrix for a ‘random walk’ or a ‘diffusion’ on the graph (see Appendix I;[59, 60]). DME consists of mapping the recording sites into an embedding space using an eigendecomposition of **P**,

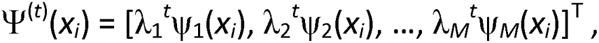

where ψ*_j_* are the eigenvectors of **P**.

The parameter *t* corresponds to the number of steps in the diffusion process (random walk on the graph). The coordinates of the data in embedding space are scaled according to λ*_i_^t^* where λ*_i_* is the eigenvalue of the *i^t^*^h^ dimension being scaled. Thus, the value of *t* sets the spatial scale of the analysis, with higher values de-emphasizing smaller eigenvalues. Because |λ*_i_*|<1 ∀ *i*, at higher values of *t* each dimension will be scaled down (‘collapse’), with the dimension corresponding to max(|λ*_i_*|) (i.e., λ_1_) scaled the least. For *t* > 5 in this dataset, the data collapses onto the single dimension of the largest eigenvalue. Because we wished to explore the structure of the data over multiple dimensions, we restricted our analyses to smaller values of *t*. Here, we present data for *t* = 1.

DME can be implemented alternatively based on a symmetric version of diffusion matrix **P_symm_** = **D**^-0.5^**KD**^-0.5^. Basing DME on **P_symm_** has the advantage that the eigenvectors of **P_symm_** form an orthogonal basis set (unlike the eigenvectors of **P**), providing some additional convenience mathematically that is beyond the scope of this paper[60]. Additionally, the eigenvalues of **P** and **P_symm_** are identical.

In two sets of analyses presented here, pairs of embeddings were compared to each other: in the analysis of lateralization of speech and language networks, and in the comparison between iEEG and fMRI data. To do that, we used a change of basis operator to map embeddings into a common embedding space using the method described in Coifman et al 2014[60].

#### Dimensionality reduction via low rank approximations to **P_symm_**

Diffusion map embedding offers an opportunity to reduce the dimensionality of the underlying data by considering only those dimensions that contribute importantly to the structure of the data, as manifested in the structure of the transition probability matrix **P**, or, equivalently, of the diffusion matrix **P_symm_**. We used the eigenvalue spectrum of **P_symm_** to determine its ideal low rank approximation, balancing dimensionality reduction and information loss. The basis for this is most easily understood in terms of the eigenvalue spectrum of **P**, whose spectrum is identical to that of **P_symm_**[60]. Because **P** is real and symmetric, the magnitude of the eigenvalues is identical to the singular values of **P**. The singular values tell us about the fidelity of low rank approximations to **P**. Specifically, if **P** has a set of singular values σ_1_≥ σ_1_≥…≥ σ_n_, then for any integer *k* ≥ 1,

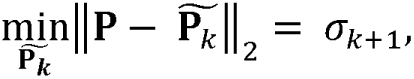

where 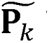 is the rank-*k* approximation to **P**. Thus, the magnitude of the eigenvalues corresponds to the fidelity of the lower dimensional approximation, and the difference in the magnitude of successive eigenvalues represents the improvement in that approximation as the dimensionality increases. The spectrum of **P** invariably has an inflection point (“elbow”), separating two sets of eigenvalues λ*_i_*: those whose magnitude decreases more quickly with increasing *i*, and those beyond the inflection point whose magnitude decreases more slowly with increasing *i*. The inflection point thus delineates the number of dimensions that are most important for approximating **P** or **P_symm_**. The inflection point *k*_infl_ was identified algorithmically[158], and the number of dimensions retained set equal to*k*_infl_ – 1.

#### Comparing distances in embedding space

The relative distance between points in embedding space provides insight into the underlying functional geometry. In several analyses presented here, two embeddings of identical sets of ROIs were compared as ROI distances within the two embeddings. After mapping to a common space and reducing dimensionality as described above, the two embeddings A and B were used to create the pairwise distance matrices À and B‘. The Pearson correlation coefficient *r* was then computed between the upper triangles (excluding the diagonal) of the corresponding elements in the distance matrices. To compare anatomical distance and distance in embedding space, inter-ROI anatomical distances were calculated for each participant by computing the centroid of each ROI in MNI space, then calculating Euclidean distances between centroids, followed by averaging distances across participants.

#### Signal to noise (SNR) characteristics

To measure the robustness of the embedding analysis to variability over time, an SNR was computed as follows. For each participant, a channel × channel **P_symm_** matrix was calculated for each 60 s segment of data. For each segment, DME analysis was applied and a channel × channel distance matrix calculated. These distance matrices were averaged across segments. The ‘signal’ of interest was defined as the variability (standard deviation) of this averaged distance matrix (ignoring the diagonals). The ‘noise’ was defined as the variability across time, estimated for each element of the distance matrix as the standard deviation across segments, then averaged across the elements of the matrix. The SNR for functional connectivity itself was computed in an analogous manner, using the original channel × channel connectivity matrix rather than the matrix of embedding distances.

#### Estimating precision in position and distances in embedding space

To obtain error estimates for both ROI locations in embedding space and embedding distance between ROIs, average ROI × ROI adjacency matrices were calculated. Our data are hierarchical/multilevel, in that we sampled participants in whom there are multiple recording sites. Nonparametric bootstrap sampling at the highest level (“cluster bootstrap”[159]; here the word “cluster” refers to the hierarchical/multilevel structure of the data, with multiple recording sites within participants, rather than algorithmic clustering) is the preferred approach for hierarchical data when groups (here, participants) are sampled and observations (here, recording sites) occur within those groups[160], with fewer necessary assumptions than multilevel (mixed-effects) modelling (e.g., subject effects are not assumed to be linear). Using this approach, participants were resampled with replacement, connectivity averaged across the bootstrapped samples, and diffusion map embedding performed for 100,000 such adjacency matrices. For locations in embedding space, these embeddings were then mapped via the change of basis procedure described above to the original group average embedding space. For each ROI, the mapped bootstrap iterations produced a cloud of locations in embedding space that were summarized by the standard deviation in each dimension. For embedding distances, no change of basis was necessary because distances are preserved across bases.

To compare the positions of STSL versus STSU relative to canonical auditory cortical ROIs (HGPM, HGAL, PT, PP, STGP, and STGM) or ROIs involved in semantic processing (STGA, MTGA, MTGP, ITGA, ITGP, TP, AGA, AGP, SMG, IFGop, IFGtr, IFGor[21, 76–78]), we calculated the average pairwise distance from STSL or STSU to each such ROI. The difference between these averages was compared to a null distribution obtained by Monte Carlo sampling of the equivalent statistic obtained by randomly exchanging STSL/STSU labels by participant. The specific comparisons performed were chosen *a priori* to constrain the number of possible hypotheses to test; pairwise comparisons of all possible ROI pairs (let alone comparisons of all higher-order groupings) would not have had sufficient statistical power under appropriate corrections for multiple comparisons. Though different choices could have been made for inclusion in the “semantic processing” category, exchanging one or two of these ROIs would not strongly influence the average distance in a group of twelve ROIs.

#### Hierarchical clustering

Agglomerative hierarchical clustering was done using the *linkage* function in MATLAB, with Euclidean distance as the distance metric and Ward’s linkage (minimum variance algorithm) as the linkage method. The ordering of ROIs along the horizontal axis in the dendrogram was determined using the *optimalleaforder* function in MATLAB, with the optimization criterion set to ‘group’.

Non-parametric bootstrapping at the participant level, as described above, was used to evaluate the robustness of clustering results both overall and at the level of individual clusters. We compared the original cluster results obtained with the full dataset to the result obtained with each bootstrap sample, then summarized those results across iterations.

Overall stability of the cluster results was evaluated using the median normalized Fowlkes-Mallows index[83] across cluster bootstrap iterations, noted as *B_k_* for *k* clusters (see Appendix II). Normalizing to the expected value *E*(*B_k_*) (see Appendix II) results in an index where 0 represents average random (chance) clustering and 1 represents perfectly identical clustering.

Cluster-wise stability was calculated by the membership of each cluster at each iteration to the corresponding cluster obtained with the full dataset using the maximum Jaccard coefficient for each reference cluster[84]. The Jaccard coefficient varies from 0 (no overlap in cluster membership) to 1 (identical membership) and is defined as the ratio of the size of the set containing intersection of the two clusters divided by the size of the set containing their union. We then subtracted from this coefficient a bias estimate calculated by randomly permuting the cluster assignments on each bootstrap iteration.

#### Auditory responsiveness

In a subset of 37 participants, auditory responsiveness was evaluated as percentage of sites within each ROI that exhibited high gamma responses to monosyllabic word stimuli. The stimuli were monosyllabic words (”cat”, “dog”, “five”, “ten”, “red”, “white”), obtained from TIMIT (https://doi.org/10.35111/17gk-bn40) and LibriVox (http://librivox.org/) databases. The words were presented in semantic categorization (animals and numbers target categories) and tone target detection tasks as described previously [23, 85–87]. A total of 20 unique exemplars of each word were presented in each task: 14 spoken by different male and 6 by different female speakers. The stimuli were delivered via insert earphones (ER4B, Etymotic Research, Elk Grove Village, IL) integrated into custom-fit earmolds. All stimuli had a duration of 300 ms, were root-mean-square amplitude-normalized and were delivered in random order. The inter-stimulus interval was chosen randomly within a Gaussian distribution (mean 2 s; SD = 10 ms). The task was to push a response button whenever the participant heard a target sound. The hand ipsilateral to the hemisphere in which the majority of electrodes were implanted was used to make the behavioral response. There was no visual component to the task, and the participants did not receive any specific instructions other than to respond to target auditory stimuli by pressing a button. Mean high gamma (70-110 Hz) power within early (50 to 350 ms) and late (350 to 650 ms) poststimulus time windows was compared with that in a prestimulus window (-200 to -100 ms). Significance of high gamma responses was established at a*α* = 0.05 level using one-tailed Mann-Whitney *U* tests with false discovery rate correction.

#### Comparing language dominant/non-dominant hemispheres

To test for differences in functional geometry between language dominant and non-dominant hemispheres, two measures were considered: differences in the location of individual ROIs in embedding space, and different pairwise distances between ROIs in embedding space. To calculate differences in location of individual ROIs, dominant/non-dominant average embeddings were mapped to a common space (from an embedding using the average across all participants regardless of language dominance) using the change of basis operator. The language-dominant location difference for a specific ROI was calculated as the Euclidean distance between the two locations of each ROI in this common space. To examine whether there was a consistent relationship between hemispheric asymmetry in a given ROI’s location in embedding space and the percentage of either early or late auditory responsive sites within that ROI, two-tailed Spearman’s rank tests were used. To calculate differences in pairwise distances between ROIs, Euclidean distances were calculated in embedding space for each hemisphere and then subtracted to obtain a difference matrix. To determine whether the differences in location or pairwise distances were larger than expected by chance, random permutations of the dominant/non-dominant labels were used to generate empirical null distributions. Since this approach produces a*p*-value for every pair of connections, *p*-values were adjusted using FDR to account for multiple comparisons.

#### Analyses of fMRI connectivity in embedding space

Two sets of analyses were performed using fMRI data. First, iEEG and fMRI data were compared in embedding space. In this analysis, connectivity based on RS-fMRI data from voxels located at electrode recording sites was compare with the corresponding connectivity matrix derived from iEEG data. The embedding analysis was applied to the two connectivity matrices, all pairwise inter-ROI distances computed, and iEEG and fMRI data compared using the correlation of the pairwise ROI distances. The second analysis was to compare embeddings derived from all ROIs in the RS-fMRI scans to those derived from just ROIs sampled with iEEG electrodes. Here, ROI × ROI connectivity matrices were computed for all ROIs, then embeddings created from the full matrices or from matrices containing just rows and columns corresponding to the ROIs sampled with iEEG.

## Data and code availability

Software and data used to generate figures are freely available at https://zenodo.org/record/7846505 or DOI 10.5281/zenodo.7846505. Complete data set is available via a request to the Authors pending establishment of a formal data sharing agreement and submission of a formal project outline. Please contact Bryan Krause for details.

## Appendix I: Diffusion Map Embedding

In the framework of DME, we consider a space *X* that is the set of *N* recording sites. We compute the similarity between those sites based on the time varying signals recorded at each site, defining similarity *k*(*x_i_,x_j_*) as the cosine similarity between functional connectivity of nodes *x_i_*and *x_j_*.

Define the matrix **K** whose *i*,*j*^th^ element is *k*(*x_i_,x_j_*). *k*(*x_i_,x_j_*) is required to be symmetric, i.e., *k*(*x_i_,x_j_*) = *k*(*x_j_,x_i_*), and positivity preserving, i.e. *k*(*x_i_,x_j_*) > 0 for all [*i,j*], to allow for spectral analysis of a normalized version of **K**.

From *X* and **K** we can construct a weighted graph Γ in which the vertices are the nodes and the edge weights are *k*(*x_i_,x_j_*). We take random walks on the graph at time steps *t* = 1, 2,…, jumping from node *x_i_* to node *x_j_* at each time step, with the (stochastic) decision as to which node should be visited next depending on *k*(*x_i_,x_j_*).

Define

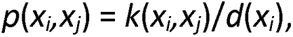

where

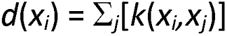

is the degree of node *x_i_*. Normalizing *k*(*x_i_,x_j_*) in this way allows us to interpret it as the probability *p*(*x_i_,x_j_*) that we’ll jump from vertex x_i_ to vertex *x_j_* in a single time step of our random walk.

If we consider a single time step, we only capture the structure in *X* on a very local scale, since we can only jump between vertices that are directly connected. As we run the random walk forward in time, we begin to explore more of our neighborhood, and we begin to explore other neighborhoods as well. Two vertices *x_i_* and *x_j_*that have similar connectivity to the rest of the network have a high probability of being connected during these longer walks because they themselves are connected to similar groups of vertices, and so there are many possible paths between *x_i_* and *x_j_*.

The diffusion operator (matrix) **P** = [*p*(*x_i_,x_j_*)] describes how signals diffuse from node to node in the graph. If **v** is a *N*×1 vector (i.e., a value assigned to each vertex, for example representing an input to each node), then **P** describes what will happen to that input as time goes on.

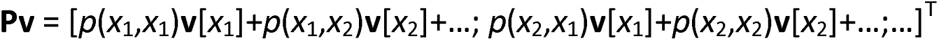

If, for example, all the nodes were insular, with *p*(*x*_i_,*x*_i_)=1 for all *i*, and otherwise *p*(*x_i_*,*x_j_*)=0, **Pv** = **v**, i.e., no diffusion occurs. If the probabilities are more distributed, **Pv** would reveal how much signals diffuse out from each node given the starting condition of **v**. Importantly, **P**^k^**v** would reveal what that distribution looks like after *k* time steps.

The eigenvector expansion of **P** based on its eigenvectors ψ*_j_* and eigenvalues, λ*_j_*, *j* = 1…*N*, is a natural method for uncovering structure in **P** because each eigenvector of **P** is a dimension along which relevant organizational features emerge. That is, clusters of related points (communities) tend to be distinct and ordered along these dimensions. In fact, we could preserve a lot of information about **P** by keeping just a subset of *M* of these vectors and discarding the rest. The information we want to preserve in the context of diffusion map embedding is the functional distance between the data at two nodes given *t* time steps to meander through the graph. We can define the diffusion map

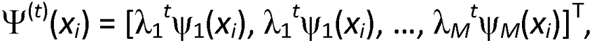

which maps each point *x* in *X* to a point in an embedding space of dimension *M* ≤ N. In this space, the diffusion distance *D*, which is the Euclidean distance between points, is the difference in the probability distributions linking *x_i_* to the rest of the network and *x_j_* to the rest of the network:

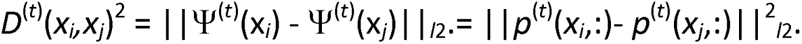

To compare embeddings across groups of participants, or modalities of measurements, it is necessary to map embeddings to a common space. To do so, consider two sets of dataα and β, and the data spaces *X*_α_ and *X*_β_. The problem is that *X*_α_ and *X*_β_ are different spaces with different kernels *k*_α_ and *k*_β_. This means that the eigenvectors for **P**_α_ and **P**_β_ will be different, and data projected into a space defined by some subset of the eigenvectors cannot be compared directly. The solution is to apply a change of basis operator to one set of the eigenvectors to get the data into the same embedding space[60]:

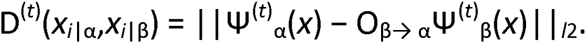

Where the change of basis operator O_β→α_ is defined as

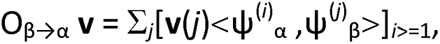

Where <ψ^(*i*)^_α_, ψ^(*j*)^_β_> is the inner product of ψ^(*i*)^_α_ and ψ^(*j*)^_β_.

## Appendix II: Fowkles-Mallows Index

The content of this appendix uses the functions and notation of Fowlkes and Mallows, 1983 {Fowlkes, 1983 #9946} with minor adjustments. Their index, denoted *B_k_*, where *k* is the number of clusters, represents the similarity of clusterings, independent of cluster ordering, as a value between 0 and 1, where 1 represents identical clustering. *B_k_* is given from the following equations:

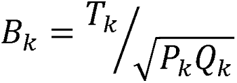

where

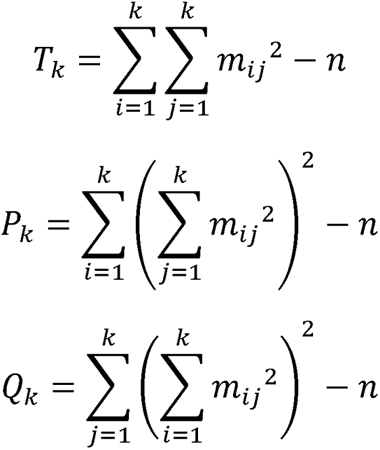

…and M = [m_ij_] is a matrix with k rows and k columns where each m_ij_ represents the number of objects in cluster i from one clustering and cluster j from the other clustering. n is the total number of objects clustered. In the case of random assignment to clusters of the sizes observed, however, *B_k_* is biased (not zero), and the expected value for this bias is given by:

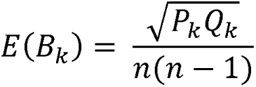

We normalized by this bias value to give a stability index that averages 0 for chance assignment to clusters (with this normalization, values less than 0 are theoretically possible) and 1 for perfect concordance:

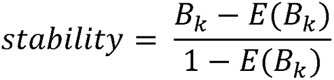

## Acknowledgements

This work was supported by the National Institutes of Health (grant numbers R01-DC04290, R01-GM109086). We are grateful to Jess Banks, Alex Billig, Haiming Chen, Phillip Gander, Christopher Garcia, Matthew Howard, Ariane Rhone, and Matthew Sutterer for help with data collection, analysis, and comments on the manuscript.

## REFERENCES CITED

1. Biswal BB, Mennes M, Zuo XN, Gohel S, Kelly C, Smith SM, et al. Toward discovery science of human brain function. Proc Natl Acad Sci U S A. 2010;107(10):4734–9. Epub 2010/02/24. doi: 10.1073/pnas.0911855107. PubMed PMID: 20176931; PubMed Central PMCID: PMCPMC2842060.

2. Yeo BT, Krienen FM, Sepulcre J, Sabuncu MR, Lashkari D, Hollinshead M, et al. The organization of the human cerebral cortex estimated by intrinsic functional connectivity. J Neurophysiol. 2011;106(3):1125–65. Epub 2011/06/10. doi: 10.1152/jn.00338.2011. PubMed PMID: 21653723; PubMed Central PMCID: PMCPMC3174820.

3. Smith SM, Fox PT, Miller KL, Glahn DC, Fox PM, Mackay CE, et al. Correspondence of the brain’s functional architecture during activation and rest. Proc Natl Acad Sci U S A. 2009;106(31):13040–5. Epub 2009/07/22. doi: 10.1073/pnas.0905267106. PubMed PMID: 19620724; PubMed Central PMCID: PMCPMC2722273.

4. Scott SK. The neurobiology of speech perception and production--can functional imaging tell us anything we did not already know? J Commun Disord. 2012;45(6):419–25. Epub 2012/07/31. doi: 10.1016/j.jcomdis.2012.06.007. PubMed PMID: 22840926.

5. Woods DL, Alain C. Functional imaging of human auditory cortex. Curr Opin Otolaryngol Head Neck Surg. 2009;17(5):407–11. Epub 2009/07/28. doi: 10.1097/MOO.0b013e3283303330. PubMed PMID: 19633556.

6. Jackson RL, Bajada CJ, Rice GE, Cloutman LL, Lambon Ralph MA. An emergent functional parcellation of the temporal cortex. Neuroimage. 2018;170:385–99. Epub 2017/04/19. doi: 10.1016/j.neuroimage.2017.04.024. PubMed PMID: 28419851.

7. Wang J, Tao A, Anderson WS, Madsen JR, Kreiman G. Mesoscopic physiological interactions in the human brain reveal small-world properties. Cell Rep. 2021;36(8):109585. Epub 2021/08/26. doi: 10.1016/j.celrep.2021.109585. PubMed PMID: 34433053; PubMed Central PMCID: PMCPMC8457376.

8. Ko AL, Weaver KE, Hakimian S, Ojemann JG. Identifying functional networks using endogenous connectivity in gamma band electrocorticography. Brain Connect. 2013;3(5):491–502. Epub 2013/07/25. doi: 10.1089/brain.2013.0157. PubMed PMID: 23879617; PubMed Central PMCID: PMCPMC3796331.

9. Zhang Y, Ding Y, Huang J, Zhou W, Ling Z, Hong B, et al. Hierarchical cortical networks of “voice patches” for processing voices in human brain. Proc Natl Acad Sci U S A. 2021;118(52). Epub 2021/12/22. doi: 10.1073/pnas.2113887118. PubMed PMID: 34930846; PubMed Central PMCID: PMCPMC8719861.

10. de Pasquale F, Della Penna S, Snyder AZ, Marzetti L, Pizzella V, Romani GL, et al. A cortical core for dynamic integration of functional networks in the resting human brain. Neuron. 2012;74(4):753–64. Epub 2012/05/29. doi: 10.1016/j.neuron.2012.03.031. PubMed PMID: 22632732; PubMed Central PMCID: PMCPMC3361697.

11. Vezoli J, Vinck M, Bosman CA, Bastos AM, Lewis CM, Kennedy H, et al. Brain rhythms define distinct interaction networks with differential dependence on anatomy. Neuron. 2021;109(23):3862–78.e5. Epub 2021/10/22. doi: 10.1016/j.neuron.2021.09.052. PubMed PMID: 34672985; PubMed Central PMCID: PMCPMC8639786.

12. Barzegaran E, Plomp G. Four concurrent feedforward and feedback networks with different roles in the visual cortical hierarchy. PLoS Biol. 2022;20(2):e3001534. Epub 2022/02/11. doi: 10.1371/journal.pbio.3001534. PubMed PMID: 35143472; PubMed Central PMCID: PMCPMC8865670.

13. Richter CG, Thompson WH, Bosman CA, Fries P. Top-Down Beta Enhances Bottom-Up Gamma. J Neurosci. 2017;37(28):6698–711. Epub 2017/06/09. doi: 10.1523/jneurosci.3771-16.2017. PubMed PMID: 28592697; PubMed Central PMCID: PMCPMC5508256.

14. Bastos AM, Vezoli J, Bosman CA, Schoffelen JM, Oostenveld R, Dowdall JR, et al. Visual areas exert feedforward and feedback influences through distinct frequency channels. Neuron. 2015;85(2):390–401. doi: 10.1016/j.neuron.2014.12.018. PubMed PMID: 25556836.

15. Michalareas G, Vezoli J, van Pelt S, Schoffelen JM, Kennedy H, Fries P. Alpha-Beta and Gamma Rhythms Subserve Feedback and Feedforward Influences among Human Visual Cortical Areas. Neuron. 2016;89(2):384–97. Epub 2016/01/19. doi: 10.1016/j.neuron.2015.12.018. PubMed PMID: 26777277; PubMed Central PMCID: PMCPMC4871751.

16. Fontolan L, Morillon B, Liegeois-Chauvel C, Giraud AL. The contribution of frequency-specific activity to hierarchical information processing in the human auditory cortex. Nat Commun. 2014;5:4694. doi: 10.1038/ncomms5694. PubMed PMID: 25178489; PubMed Central PMCID: PMCPMC4164774.

17. Baroni F, Morillon B, Trébuchon A, Liégeois-Chauvel C, Olasagasti I, Giraud AL. Converging intracortical signatures of two separated processing timescales in human early auditory cortex. Neuroimage. 2020;218:116882. Epub 2020/05/23. doi: 10.1016/j.neuroimage.2020.116882. PubMed PMID: 32439539.

18. Chalas N, Omigie D, Poeppel D, van Wassenhove V. Hierarchically nested networks optimize the analysis of audiovisual speech. iScience. 2023;26(3):106257. Epub 2023/03/14. doi: 10.1016/j.isci.2023.106257. PubMed PMID: 36909667; PubMed Central PMCID: PMCPMC9993032.

19. Hayat H, Marmelshtein A, Krom AJ, Sela Y, Tankus A, Strauss I, et al. Reduced neural feedback signaling despite robust neuron and gamma auditory responses during human sleep. Nat Neurosci. 2022;25(7):935–43. Epub 2022/07/12. doi: 10.1038/s41593-022-01107-4. PubMed PMID: 35817847; PubMed Central PMCID: PMCPMC9276533.

20. Visser M, Jefferies E, Lambon Ralph MA. Semantic processing in the anterior temporal lobes: a meta-analysis of the functional neuroimaging literature. J Cogn Neurosci. 2010;22(6):1083–94. Epub 2009/07/09. doi: 10.1162/jocn.2009.21309. PubMed PMID: 19583477.

21. Ralph MA, Jefferies E, Patterson K, Rogers TT. The neural and computational bases of semantic cognition. Nat Rev Neurosci. 2017;18(1):42–55. Epub 2016/11/25. doi: 10.1038/nrn.2016.150. PubMed PMID: 27881854.

22. Abrams DA, Kochalka J, Bhide S, Ryali S, Menon V. Intrinsic functional architecture of the human speech processing network. Cortex. 2020;129:41–56. Epub 2020/05/20. doi: 10.1016/j.cortex.2020.03.013. PubMed PMID: 32428761.

23. Nourski KV, Steinschneider M, Rhone AE, Kovach CK, Banks MI, Krause BM, et al. Electrophysiology of the Human Superior Temporal Sulcus during Speech Processing. Cereb Cortex. 2021;31(2):1131–48. Epub 2020/10/17. doi: 10.1093/cercor/bhaa281. PubMed PMID: 33063098; PubMed Central PMCID: PMCPMC7786351.

24. Langs G, Golland P, Tie Y, Rigolo L, Golby AJ. Functional Geometry Alignment and Localization of Brain Areas. Adv Neural Inf Process Syst. 2010;1:1225–33. Epub 2010/01/01. PubMed PMID: 24808719; PubMed Central PMCID: PMCPMC4010233.

25. Margulies DS, Ghosh SS, Goulas A, Falkiewicz M, Huntenburg JM, Langs G, et al. Situating the default-mode network along a principal gradient of macroscale cortical organization. Proc Natl Acad Sci U S A. 2016;113(44):12574–9. Epub 2016/11/03. doi: 10.1073/pnas.1608282113. PubMed PMID: 27791099; PubMed Central PMCID: PMCPMC5098630.

26. Huang Z, Mashour GA, Hudetz AG. Functional geometry of the cortex encodes dimensions of consciousness. Nat Commun. 2023;14(1):72. Epub 2023/01/06. doi: 10.1038/s41467-022-35764-7. PubMed PMID: 36604428; PubMed Central PMCID: PMCPMC9814511.

27. Wilson SM, Bautista A, McCarron A. Convergence of spoken and written language processing in the superior temporal sulcus. Neuroimage. 2018;171:62–74. Epub 2017/12/27. doi: 10.1016/j.neuroimage.2017.12.068. PubMed PMID: 29277646; PubMed Central PMCID: PMCPMC5857434.

28. Forseth KJ, Hickok G, Rollo PS, Tandon N. Language prediction mechanisms in human auditory cortex. Nat Commun. 2020;11(1):5240. Epub 2020/10/18. doi: 10.1038/s41467-020-19010-6. PubMed PMID: 33067457; PubMed Central PMCID: PMCPMC7567874.

29. Hamilton LS, Oganian Y, Hall J, Chang EF. Parallel and distributed encoding of speech across human auditory cortex. Cell. 2021;184(18):4626–39 e13. Epub 2021/08/20. doi: 10.1016/j.cell.2021.07.019. PubMed PMID: 34411517; PubMed Central PMCID: PMCPMC8456481.

30. Friederici AD, Meyer M, von Cramon DY. Auditory language comprehension: an event-related fMRI study on the processing of syntactic and lexical information. Brain Lang. 2000;75(3):289–300. Epub 2001/06/02. PubMed PMID: 11386224.

31. Angulo-Perkins A, Aube W, Peretz I, Barrios FA, Armony JL, Concha L. Music listening engages specific cortical regions within the temporal lobes: differences between musicians and non-musicians. Cortex. 2014;59:126–37. Epub 2014/09/01. doi: 10.1016/j.cortex.2014.07.013. PubMed PMID: 25173956.

32. Zhang Y, Zhou W, Wang S, Zhou Q, Wang H, Zhang B, et al. The Roles of Subdivisions of Human Insula in Emotion Perception and Auditory Processing. Cereb Cortex. 2019;29(2):517–28. Epub 2018/01/18. doi: 10.1093/cercor/bhx334. PubMed PMID: 29342237.

33. Price CJ. A review and synthesis of the first 20 years of PET and fMRI studies of heard speech, spoken language and reading. Neuroimage. 2012;62(2):816–47. Epub 2012/05/16. doi: 10.1016/j.neuroimage.2012.04.062. PubMed PMID: 22584224; PubMed Central PMCID: PMCPMC3398395.

34. Hickok G. The functional neuroanatomy of language. Phys Life Rev. 2009;6(3):121–43. Epub 2010/02/18. doi: 10.1016/j.plrev.2009.06.001. PubMed PMID: 20161054; PubMed Central PMCID: PMCPMC2747108.

35. Beauchamp MS. The social mysteries of the superior temporal sulcus. Trends Cogn Sci. 2015;19(9):489–90. Epub 2015/07/26. doi: 10.1016/j.tics.2015.07.002. PubMed PMID: 26208834; PubMed Central PMCID: PMCPMC4556565.

36. Chang EF, Raygor KP, Berger MS. Contemporary model of language organization: an overview for neurosurgeons. J Neurosurg. 2015;122(2):250–61. Epub 2014/11/26. doi: 10.3171/2014.10.JNS132647. PubMed PMID: 25423277.

37. Venezia JH, Vaden KI, Jr., Rong F, Maddox D, Saberi K, Hickok G. Auditory, Visual and Audiovisual Speech Processing Streams in Superior Temporal Sulcus. Front Hum Neurosci. 2017;11:174. Epub 2017/04/26. doi: 10.3389/fnhum.2017.00174. PubMed PMID: 28439236; PubMed Central PMCID: PMCPMC5383672.

38. Rauschecker JP, Scott SK. Maps and streams in the auditory cortex: nonhuman primates illuminate human speech processing. Nat Neurosci. 2009;12(6):718–24. doi: 10.1038/nn.2331. PubMed PMID: 19471271; PubMed Central PMCID: PMCPMC2846110.

39. Hickok G, Poeppel D. The cortical organization of speech processing. Nat Rev Neurosci. 2007;8(5):393–402. Epub 2007/04/14. doi: 10.1038/nrn2113. PubMed PMID: 17431404.

40. Friederici AD. The cortical language circuit: from auditory perception to sentence comprehension. Trends Cogn Sci. 2012;16(5):262–8. Epub 2012/04/21. doi: 10.1016/j.tics.2012.04.001. PubMed PMID: 22516238.

41. Cloutman LL. Interaction between dorsal and ventral processing streams: where, when and how? Brain Lang. 2013;127(2):251–63. Epub 2012/09/13. doi: 10.1016/j.bandl.2012.08.003. PubMed PMID: 22968092.

42. Hickok G, Poeppel D. Neural basis of speech perception. Handb Clin Neurol. 2015;129:149–60. Epub 2015/03/03. doi: 10.1016/B978-0-444-62630-1.00008-1. PubMed PMID: 25726267.

43. Rauschecker JP. Where, When, and How: Are they all sensorimotor? Towards a unified view of the dorsal pathway in vision and audition. Cortex. 2018;98:262–8. Epub 2017/12/01. doi: 10.1016/j.cortex.2017.10.020. PubMed PMID: 29183630; PubMed Central PMCID: PMCPMC5771843.

44. Saur D, Kreher BW, Schnell S, Kummerer D, Kellmeyer P, Vry MS, et al. Ventral and dorsal pathways for language. Proc Natl Acad Sci U S A. 2008;105(46):18035–40. Epub 2008/11/14. doi: 10.1073/pnas.0805234105. PubMed PMID: 19004769; PubMed Central PMCID: PMCPMC2584675.

45. Munoz-Lopez MM, Mohedano-Moriano A, Insausti R. Anatomical pathways for auditory memory in primates. Front Neuroanat. 2010;4:129. Epub 2010/10/27. doi: 10.3389/fnana.2010.00129. PubMed PMID: 20976037; PubMed Central PMCID: PMCPMC2957958.

46. Kraus KS, Canlon B. Neuronal connectivity and interactions between the auditory and limbic systems. Effects of noise and tinnitus. Hear Res. 2012;288(1-2):34–46. Epub 2012/03/24. doi: 10.1016/j.heares.2012.02.009. PubMed PMID: 22440225.

47. Husain FT, Schmidt SA. Using resting state functional connectivity to unravel networks of tinnitus. Hear Res. 2014;307:153–62. Epub 2013/07/31. doi: 10.1016/j.heares.2013.07.010. PubMed PMID: 23895873.

48. Kumar S, Joseph S, Gander PE, Barascud N, Halpern AR, Griffiths TD. A Brain System for Auditory Working Memory. J Neurosci. 2016;36(16):4492–505. Epub 2016/04/22. doi: 10.1523/jneurosci.4341-14.2016. PubMed PMID: 27098693; PubMed Central PMCID: PMCPMC4837683.

49. Kumar S, Gander PE, Berger JI, Billig AJ, Nourski KV, Oya H, et al. Oscillatory correlates of auditory working memory examined with human electrocorticography. Neuropsychologia. 2021;150:107691. Epub 2020/11/24. doi: 10.1016/j.neuropsychologia.2020.107691. PubMed PMID: 33227284; PubMed Central PMCID: PMCPMC7884909.

50. Geschwind N. The organization of language and the brain. Science. 1970;170(3961):940-4. Epub 1970/11/27. doi: 10.1126/science.170.3961.940. PubMed PMID: 5475022.

51. Hagoort P. The neurobiology of language beyond single-word processing. Science. 2019;366(6461):55-8. Epub 2019/10/12. doi: 10.1126/science.aax0289. PubMed PMID: 31604301.

52. de Heer WA, Huth AG, Griffiths TL, Gallant JL, Theunissen FE. The Hierarchical Cortical Organization of Human Speech Processing. J Neurosci. 2017;37(27):6539–57. Epub 2017/06/08. doi: 10.1523/jneurosci.3267-16.2017. PubMed PMID: 28588065; PubMed Central PMCID: PMCPMC5511884.

53. Binder JR, Frost JA, Hammeke TA, Bellgowan PS, Springer JA, Kaufman JN, et al. Human temporal lobe activation by speech and nonspeech sounds. Cereb Cortex. 2000;10(5):512–28. Epub 2000/06/10. doi: 10.1093/cercor/10.5.512. PubMed PMID: 10847601.

54. Cogan GB, Thesen T, Carlson C, Doyle W, Devinsky O, Pesaran B. Sensory-motor transformations for speech occur bilaterally. Nature. 2014;507(7490):94-8. Epub 2014/01/17. doi: 10.1038/nature12935. PubMed PMID: 24429520; PubMed Central PMCID: PMCPMC4000028.

55. McGettigan C, Scott SK. Cortical asymmetries in speech perception: what’s wrong, what’s right and what’s left? Trends Cogn Sci. 2012;16(5):269–76. Epub 2012/04/24. doi: 10.1016/j.tics.2012.04.006. PubMed PMID: 22521208; PubMed Central PMCID: PMCPMC4083255.

56. Turkeltaub PE, Coslett HB. Localization of sublexical speech perception components. Brain Lang. 2010;114(1):1–15. Epub 2010/04/24. doi: 10.1016/j.bandl.2010.03.008. PubMed PMID: 20413149; PubMed Central PMCID: PMCPMC2914564.

57. Leaver AM, Rauschecker JP. Cortical representation of natural complex sounds: effects of acoustic features and auditory object category. J Neurosci. 2010;30(22):7604–12. Epub 2010/06/04. doi: 10.1523/jneurosci.0296-10.2010. PubMed PMID: 20519535; PubMed Central PMCID: PMCPMC2930617.

58. Eisner F, McGettigan C, Faulkner A, Rosen S, Scott SK. Inferior frontal gyrus activation predicts individual differences in perceptual learning of cochlear-implant simulations. J Neurosci. 2010;30(21):7179–86. Epub 2010/05/28. doi: 10.1523/JNEUROSCI.4040-09.2010. PubMed PMID: 20505085; PubMed Central PMCID: PMCPMC2883443.

59. Coifman RR, Lafon S, Lee AB, Maggioni M, Nadler B, Warner F, et al. Geometric diffusions as a tool for harmonic analysis and structure definition of data: diffusion maps. Proc Natl Acad Sci U S A. 2005;102(21):7426–31. Epub 2005/05/19. doi: 10.1073/pnas.0500334102. PubMed PMID: 15899970; PubMed Central PMCID: PMCPMC1140422.

60. Coifman RR, Hirn MJ. Diffusion maps for changing data. Applied and Computational Harmonic Analysis. 2014;36(1):79–107. doi: https://doi.org/10.1016/j.acha.2013.03.001.

61. Lafon S, Lee AB. Diffusion maps and coarse-graining: A unified framework for dimensionality reduction, graph partitioning, and data set parameterization. IEEE Trans Pattern Anal Mach Intell. 2006;28(9):1393–403. Epub 2006/08/26. doi: 10.1109/TPAMI.2006.184. PubMed PMID: 16929727.

62. Banks MI, Krause BM, Endemann CM, Campbell DI, Kovach CK, Dyken ME, et al. Cortical functional connectivity indexes arousal state during sleep and anesthesia. Neuroimage. 2020;211:116627. Epub 2020/02/12. doi: 10.1016/j.neuroimage.2020.116627. PubMed PMID: 32045640; PubMed Central PMCID: PMCPMC7117963.

63. Nourski KV, Steinschneider M, Rhone AE, Kawasaki H, Howard MA, 3rd, Banks MI. Auditory Predictive Coding across Awareness States under Anesthesia: An Intracranial Electrophysiology Study. J Neurosci. 2018;38(39):8441–52. Epub 2018/08/22. doi: 10.1523/JNEUROSCI.0967-18.2018. PubMed PMID: 30126970; PubMed Central PMCID: PMCPMC6158689.

64. Nourski KV, Steinschneider M, Rhone AE, Kawasaki H, Howard MA, 3rd, Banks MI. Processing of auditory novelty across the cortical hierarchy: An intracranial electrophysiology study. Neuroimage. 2018;183:412–24. Epub 2018/08/17. doi: 10.1016/j.neuroimage.2018.08.027. PubMed PMID: 30114466; PubMed Central PMCID: PMCPMC6207077

65. Nourski KV, Steinschneider M, Rhone AE, Krause BM, Kawasaki H, Banks MI. Cortical responses to auditory novelty across task conditions: An intracranial electrophysiology study. Hear Res. 2021;399:107911. Epub 2020/02/23. doi: 10.1016/j.heares.2020.107911. PubMed PMID: 32081413; PubMed Central PMCID: PMCPMC7417283.

66. Nourski KV, Steinschneider M, Rhone AE, Krause BM, Mueller RN, Kawasaki H, et al. Cortical Responses to Vowel Sequences in Awake and Anesthetized States: A Human Intracranial Electrophysiology Study. Cereb Cortex. 2021;31(12):5435–48. Epub 2021/06/13. doi: 10.1093/cercor/bhab168. PubMed PMID: 34117741; PubMed Central PMCID: PMCPMC8568007.

67. Nourski KV, Steinschneider M, Rhone AE, Mueller RN, Kawasaki H, Banks MI. Arousal State-Dependence of Interactions Between Short- and Long-Term Auditory Novelty Responses in Human Subjects. Front Hum Neurosci. 2021;15:737230. Epub 2021/10/19. doi: 10.3389/fnhum.2021.737230. PubMed PMID: 34658820; PubMed Central PMCID: PMCPMC8517406.

68. Hipp JF, Hawellek DJ, Corbetta M, Siegel M, Engel AK. Large-scale cortical correlation structure of spontaneous oscillatory activity. Nat Neurosci. 2012;15(6):884–90. Epub 2012/05/09. doi: 10.1038/nn.3101. PubMed PMID: 22561454; PubMed Central PMCID: PMCPMC3861400.

69. Ercsey-Ravasz M, Markov NT, Lamy C, Van Essen DC, Knoblauch K, Toroczkai Z, et al. A predictive network model of cerebral cortical connectivity based on a distance rule. Neuron. 2013;80(1):184–97. Epub 2013/10/08. doi: 10.1016/j.neuron.2013.07.036. PubMed PMID: 24094111; PubMed Central PMCID: PMCPMC3954498.

70. Horvát S, Gămănuț R, Ercsey-Ravasz M, Magrou L, Gămănuț B, Van Essen DC, et al. Spatial Embedding and Wiring Cost Constrain the Functional Layout of the Cortical Network of Rodents and Primates. PLoS Biol. 2016;14(7):e1002512. Epub 2016/07/22. doi: 10.1371/journal.pbio.1002512. PubMed PMID: 27441598; PubMed Central PMCID: PMCPMC4956175.

71. Theodoni P, Majka P, Reser DH, Wójcik DK, Rosa MGP, Wang XJ. Structural Attributes and Principles of the Neocortical Connectome in the Marmoset Monkey. Cereb Cortex. 2021;32(1):15–28. Epub 2021/07/19. doi: 10.1093/cercor/bhab191. PubMed PMID: 34274966; PubMed Central PMCID: PMCPMC8634603.

72. Remedios R, Logothetis NK, Kayser C. An auditory region in the primate insular cortex responding preferentially to vocal communication sounds. J Neurosci. 2009;29(4):1034–45. Epub 2009/01/30. doi: 10.1523/JNEUROSCI.4089-08.2009. PubMed PMID: 19176812; PubMed Central PMCID: PMCPMC6665141.

73. Steinschneider M, Nourski KV, Fishman YI. Representation of speech in human auditory cortex: is it special? Hear Res. 2013;305:57–73. doi: 10.1016/j.heares.2013.05.013. PubMed PMID: 23792076; PubMed Central PMCID: PMCPMC3818517.

74. Craig AD. Interoception: the sense of the physiological condition of the body. Curr Opin Neurobiol. 2003;13(4):500–5. Epub 2003/09/11. doi: 10.1016/s0959-4388(03)00090-4. PubMed PMID: 12965300.

75. Kuehn E, Mueller K, Lohmann G, Schuetz-Bosbach S. Interoceptive awareness changes the posterior insula functional connectivity profile. Brain Struct Funct. 2016;221(3):1555–71. Epub 2015/01/24. doi: 10.1007/s00429-015-0989-8. PubMed PMID: 25613901.

76. Binder JR, Desai RH, Graves WW, Conant LL. Where is the semantic system? A critical review and meta-analysis of 120 functional neuroimaging studies. Cereb Cortex. 2009;19(12):2767–96. Epub 2009/03/31. doi: 10.1093/cercor/bhp055. PubMed PMID: 19329570; PubMed Central PMCID: PMCPMC2774390.

77. Humphreys GF, Hoffman P, Visser M, Binney RJ, Lambon Ralph MA. Establishing task- and modality-dependent dissociations between the semantic and default mode networks. Proc Natl Acad Sci U S A. 2015;112(25):7857–62. Epub 2015/06/10. doi: 10.1073/pnas.1422760112. PubMed PMID: 26056304; PubMed Central PMCID: PMCPMC4485123.

78. Jackson RL, Hoffman P, Pobric G, Lambon Ralph MA. The Semantic Network at Work and Rest: Differential Connectivity of Anterior Temporal Lobe Subregions. J Neurosci. 2016;36(5):1490–501. Epub 2016/02/05. doi: 10.1523/JNEUROSCI.2999-15.2016. PubMed PMID: 26843633; PubMed Central PMCID: PMCPMC4737765.

79. Bernstein LE, Liebenthal E. Neural pathways for visual speech perception. Front Neurosci. 2014;8:386. Epub 2014/12/19. doi: 10.3389/fnins.2014.00386. PubMed PMID: 25520611; PubMed Central PMCID: PMCPMC4248808.

80. Hacker CD, Snyder AZ, Pahwa M, Corbetta M, Leuthardt EC. Frequency-specific electrophysiologic correlates of resting state fMRI networks. Neuroimage. 2017;149:446–57. Epub 2017/02/06. doi: 10.1016/j.neuroimage.2017.01.054. PubMed PMID: 28159686; PubMed Central PMCID: PMCPMC5745814.

81. Kiebel SJ, Daunizeau J, Friston KJ. A hierarchy of time-scales and the brain. PLoS Comput Biol. 2008;4(11):e1000209. Epub 2008/11/15. doi: 10.1371/journal.pcbi.1000209. PubMed PMID: 19008936; PubMed Central PMCID: PMCPMC2568860.

82. Keitel A, Gross J. Individual Human Brain Areas Can Be Identified from Their Characteristic Spectral Activation Fingerprints. PLoS Biol. 2016;14(6):e1002498. Epub 2016/06/30. doi: 10.1371/journal.pbio.1002498. PubMed PMID: 27355236; PubMed Central PMCID: PMCPMC4927181.

83. Fowlkes EB, Mallows CL. A Method for Comparing Two Hierarchical Clusterings. Journal of the American Statistical Association. 1983;78(383):553–69. doi: 10.1080/01621459.1983.10478008.

84. Hennig C. Cluster-wise assessment of cluster stability. Computational Statistics & Data Analysis. 2007;52(1):258–71. doi: https://doi.org/10.1016/j.csda.2006.11.025.

85. Nourski KV, Steinschneider M, Rhone AE, Howard Iii MA. Intracranial Electrophysiology of Auditory Selective Attention Associated with Speech Classification Tasks. Front Hum Neurosci. 2016;10:691. Epub 2017/01/26. doi: 10.3389/fnhum.2016.00691. PubMed PMID: 28119593; PubMed Central PMCID: PMCPMC5222875.

86. Steinschneider M, Nourski KV, Rhone AE, Kawasaki H, Oya H, Howard MA, 3rd. Differential activation of human core, non-core and auditory-related cortex during speech categorization tasks as revealed by intracranial recordings. Front Neurosci. 2014;8:240. doi: 10.3389/fnins.2014.00240. PubMed PMID: 25157216; PubMed Central PMCID: PMCPMC4128221.

87. Nourski KV, Steinschneider M, Rhone AE, Kovach CK, Kawasaki H, Howard MA, 3rd. Gamma Activation and Alpha Suppression within Human Auditory Cortex during a Speech Classification Task. J Neurosci. 2022;42(25):5034–46. Epub 2022/05/10. doi: 10.1523/JNEUROSCI.2187-21.2022. PubMed PMID: 35534226; PubMed Central PMCID: PMCPMC9233444.

88. Schaefer A, Kong R, Gordon EM, Laumann TO, Zuo XN, Holmes AJ, et al. Local-Global Parcellation of the Human Cerebral Cortex from Intrinsic Functional Connectivity MRI. Cereb Cortex. 2018;28(9):3095–114. Epub 2017/10/06. doi: 10.1093/cercor/bhx179. PubMed PMID: 28981612; PubMed Central PMCID: PMCPMC6095216.

89. Bullmore E, Sporns O. Complex brain networks: graph theoretical analysis of structural and functional systems. NatRevNeurosci. 2009;10(3):186–98. doi: nrn2575 [pii];10.1038/nrn2575 [doi].

90. Chapter 5 - Centrality and Hubs. In: Fornito A, Zalesky A, Bullmore ET, editors. Fundamentals of Brain Network Analysis. San Diego: Academic Press; 2016. p. 137–61.

91. Knecht S, Drager B, Deppe M, Bobe L, Lohmann H, Floel A, et al. Handedness and hemispheric language dominance in healthy humans. Brain. 2000;123 Pt 12:2512–8. Epub 2000/12/02. doi: 10.1093/brain/123.12.2512. PubMed PMID: 11099452.

92. Schirmer A, Fox PM, Grandjean D. On the spatial organization of sound processing in the human temporal lobe: a meta-analysis. Neuroimage. 2012;63(1):137–47. Epub 2012/06/27. doi: 10.1016/j.neuroimage.2012.06.025. PubMed PMID: 22732561.

93. Ardila A, Bernal B, Rosselli M. How Localized are Language Brain Areas? A Review of Brodmann Areas Involvement in Oral Language. Arch Clin Neuropsychol. 2016;31(1):112–22. Epub 2015/12/15. doi: 10.1093/arclin/acv081. PubMed PMID: 26663825.

94. Kucyi A, Schrouff J, Bickel S, Foster BL, Shine JM, Parvizi J. Intracranial Electrophysiology Reveals Reproducible Intrinsic Functional Connectivity within Human Brain Networks. J Neurosci. 2018;38(17):4230–42. Epub 2018/04/08. doi: 10.1523/jneurosci.0217-18.2018. PubMed PMID: 29626167; PubMed Central PMCID: PMCPMC5963853.

95. Hull JV, Dokovna LB, Jacokes ZJ, Torgerson CM, Irimia A, Van Horn JD. Resting-State Functional Connectivity in Autism Spectrum Disorders: A Review. Front Psychiatry. 2016;7:205. Epub 2017/01/20. doi: 10.3389/fpsyt.2016.00205. PubMed PMID: 28101064; PubMed Central PMCID: PMCPMC5209637.

96. Badhwar A, Tam A, Dansereau C, Orban P, Hoffstaedter F, Bellec P. Resting-state network dysfunction in Alzheimer’s disease: A systematic review and meta-analysis. Alzheimers Dement (Amst). 2017;8:73–85. Epub 2017/06/01. doi: 10.1016/j.dadm.2017.03.007. PubMed PMID: 28560308; PubMed Central PMCID: PMCPMC5436069.

97. Sha Z, Wager TD, Mechelli A, He Y. Common Dysfunction of Large-Scale Neurocognitive Networks Across Psychiatric Disorders. Biol Psychiatry. 2019;85(5):379–88. Epub 2019/01/08. doi: 10.1016/j.biopsych.2018.11.011. PubMed PMID: 30612699.

98. Sanders RDB, M. I.; Darracq, M.; Moran, R.; Sleigh, J.; Gosseries, O.; Bonhomme, V.; Brichant, J-F.; Rosonova, M.; Raz, A.; Tononi, G.; Massimini, M.; Laureys, S.; Boly, M. Propofol-Induced Unresponsiveness is Associated with Impaired Feedforward Connectivity in the Cortical Hierarchy. bioRxiv. 2017;213504. doi: https://doi.org/10.1101/213504.

99. Huang Z, Tarnal V, Vlisides PE, Janke EL, McKinney AM, Picton P, et al. Asymmetric neural dynamics characterize loss and recovery of consciousness. Neuroimage. 2021;236:118042. Epub 2021/04/14. doi: 10.1016/j.neuroimage.2021.118042. PubMed PMID: 33848623; PubMed Central PMCID: PMCPMC8310457.

100. Van Essen DC, Donahue C, Dierker DL, Glasser MF. Parcellations and Connectivity Patterns in Human and Macaque Cerebral Cortex. In: Kennedy H, Van Essen DC, Christen Y, editors. Micro-, Meso- and Macro-Connectomics of the Brain. Cham (CH): Springer Copyright 2016, The Author(s). 2016. p. 89–106.

101. Hackett TA, Preuss TM, Kaas JH. Architectonic identification of the core region in auditory cortex of macaques, chimpanzees, and humans. J Comp Neurol. 2001;441(3):197–222.

102. Hackett TA. Anatomic organization of the auditory cortex. Handb Clin Neurol. 2015;129:27–53. doi: 10.1016/B978-0-444-62630-1.00002-0. PubMed PMID: 25726261.

103. Woods DL, Herron TJ, Cate AD, Yund EW, Stecker GC, Rinne T, et al. Functional properties of human auditory cortical fields. Front Syst Neurosci. 2010;4:155. Epub 2010/12/17. doi: 10.3389/fnsys.2010.00155. PubMed PMID: 21160558; PubMed Central PMCID: PMCPMC3001989.

104. Barton B, Venezia JH, Saberi K, Hickok G, Brewer AA. Orthogonal acoustic dimensions define auditory field maps in human cortex. Proc Natl Acad Sci U S A. 2012;109(50):20738–43. Epub 2012/11/29. doi: 10.1073/pnas.1213381109. PubMed PMID: 23188798; PubMed Central PMCID: PMCPMC3528571.

105. Moerel M, De Martino F, Formisano E. An anatomical and functional topography of human auditory cortical areas. Front Neurosci. 2014;8:225. Epub 2014/08/15. doi: 10.3389/fnins.2014.00225. PubMed PMID: 25120426; PubMed Central PMCID: PMCPMC4114190.

106. Howard MA, Volkov IO, Mirsky R, Garell PC, Noh MD, Granner M, et al. Auditory cortex on the human posterior superior temporal gyrus. J Comp Neurol. 2000;416(1):79–92.

107. Nourski KV, Steinschneider M, Oya H, Kawasaki H, Jones RD, Howard MA. Spectral organization of the human lateral superior temporal gyrus revealed by intracranial recordings. Cereb Cortex. 2014;24(2):340–52. doi: 10.1093/cercor/bhs314. PubMed PMID: 23048019; PubMed Central PMCID: PMCPMC3888366.

108. Upadhyay J, Silver A, Knaus TA, Lindgren KA, Ducros M, Kim DS, et al. Effective and structural connectivity in the human auditory cortex. J Neurosci. 2008;28(13):3341–9. Epub 2008/03/28. doi: 10.1523/JNEUROSCI.4434-07.2008. PubMed PMID: 18367601; PubMed Central PMCID: PMCPMC6670606.

109. Zachlod D, Rüttgers B, Bludau S, Mohlberg H, Langner R, Zilles K, et al. Four new cytoarchitectonic areas surrounding the primary and early auditory cortex in human brains. Cortex. 2020;128:1–21. Epub 2020/04/17. doi: 10.1016/j.cortex.2020.02.021. PubMed PMID: 32298845.

110. Belin P, Zatorre RJ, Lafaille P, Ahad P, Pike B. Voice-selective areas in human auditory cortex. Nature. 2000;403(6767):309–12.

111. Deen B, Koldewyn K, Kanwisher N, Saxe R. Functional Organization of Social Perception and Cognition in the Superior Temporal Sulcus. Cereb Cortex. 2015;25(11):4596–609. Epub 2015/06/07. doi: 10.1093/cercor/bhv111. PubMed PMID: 26048954; PubMed Central PMCID: PMCPMC4816802.

112. Kahn I, Andrews-Hanna JR, Vincent JL, Snyder AZ, Buckner RL. Distinct cortical anatomy linked to subregions of the medial temporal lobe revealed by intrinsic functional connectivity. J Neurophysiol. 2008;100(1):129–39. Epub 2008/04/04. doi: 10.1152/jn.00077.2008. PubMed PMID: 18385483; PubMed Central PMCID: PMCPMC2493488.

113. Wang SF, Ritchey M, Libby LA, Ranganath C. Functional connectivity based parcellation of the human medial temporal lobe. Neurobiol Learn Mem. 2016;134 Pt A:123–34. Epub 2016/01/26. doi: 10.1016/j.nlm.2016.01.005. PubMed PMID: 26805590; PubMed Central PMCID: PMCPMC4955645.

114. Michelmann S, Price AR, Aubrey B, Strauss CK, Doyle WK, Friedman D, et al. Moment-by-moment tracking of naturalistic learning and its underlying hippocampo-cortical interactions. Nat Commun. 2021;12(1):5394. Epub 2021/09/15. doi: 10.1038/s41467-021-25376-y. PubMed PMID: 34518520; PubMed Central PMCID: PMCPMC8438040.

115. Rocchi F, Oya H, Balezeau F, Billig AJ, Kocsis Z, Jenison RL, et al. Common fronto-temporal effective connectivity in humans and monkeys. Neuron. 2021;109(5):852–68 e8. Epub 2021/01/23. doi: 10.1016/j.neuron.2020.12.026. PubMed PMID: 33482086; PubMed Central PMCID: PMCPMC7927917.

116. Fruhholz S, Trost W, Kotz SA. The sound of emotions-Towards a unifying neural network perspective of affective sound processing. Neurosci Biobehav Rev. 2016;68:96–110. Epub 2016/05/18. doi: 10.1016/j.neubiorev.2016.05.002. PubMed PMID: 27189782.

117. Munoz-Lopez M, Insausti R, Mohedano-Moriano A, Mishkin M, Saunders RC. Anatomical pathways for auditory memory II: information from rostral superior temporal gyrus to dorsolateral temporal pole and medial temporal cortex. Front Neurosci. 2015;9:158. Epub 2015/06/05. doi: 10.3389/fnins.2015.00158. PubMed PMID: 26041980; PubMed Central PMCID: PMCPMC4435056.

118. Olson IR, McCoy D, Klobusicky E, Ross LA. Social cognition and the anterior temporal lobes: a review and theoretical framework. Soc Cogn Affect Neurosci. 2013;8(2):123–33. Epub 2012/10/12. doi: 10.1093/scan/nss119. PubMed PMID: 23051902; PubMed Central PMCID: PMCPMC3575728.

119. Mesulam MM. Paralimbic (mesocortical) areas. Principles of behavioral and cognitive neurology. New York, NY: Oxford University Press; 2000. p. 49–54.

120. Chanes L, Barrett LF. Redefining the Role of Limbic Areas in Cortical Processing. Trends Cogn Sci. 2016;20(2):96–106. Epub 2015/12/26. doi: 10.1016/j.tics.2015.11.005. PubMed PMID: 26704857; PubMed Central PMCID: PMCPMC4780414.

121. Maller JJ, Welton T, Middione M, Callaghan FM, Rosenfeld JV, Grieve SM. Revealing the Hippocampal Connectome through Super-Resolution 1150-Direction Diffusion MRI. Sci Rep. 2019;9(1):2418. Epub 2019/02/23. doi: 10.1038/s41598-018-37905-9. PubMed PMID: 30787303; PubMed Central PMCID: PMCPMC6382767.

122. Hickok G. Computational neuroanatomy of speech production. Nat Rev Neurosci. 2012;13(2):135–45. Epub 2012/01/06. doi: 10.1038/nrn3158. PubMed PMID: 22218206; PubMed Central PMCID: PMCPMC5367153.

123. Rauschecker JP. An expanded role for the dorsal auditory pathway in sensorimotor control and integration. Hear Res. 2011;271(1-2):16–25. Epub 2010/09/21. doi: 10.1016/j.heares.2010.09.001. PubMed PMID: 20850511; PubMed Central PMCID: PMCPMC3021714.

124. Smith E, Duede S, Hanrahan S, Davis T, House P, Greger B. Seeing is believing: neural representations of visual stimuli in human auditory cortex correlate with illusory auditory perceptions. PLoS One. 2013;8(9):e73148. Epub 2013/09/12. doi: 10.1371/journal.pone.0073148. PubMed PMID: 24023823; PubMed Central PMCID: PMCPMC3762867.

125. Rolls ET, Rauschecker JP, Deco G, Huang CC, Feng J. Auditory cortical connectivity in humans. Cereb Cortex. 2022. Epub 2022/12/28. doi: 10.1093/cercor/bhac496. PubMed PMID: 36573464.

126. van den Heuvel MP, Sporns O. Network hubs in the human brain. Trends Cogn Sci. 2013;17(12):683–96. Epub 2013/11/16. doi: 10.1016/j.tics.2013.09.012. PubMed PMID: 24231140.

127. Simmons WK, Martin A. The anterior temporal lobes and the functional architecture of semantic memory. J Int Neuropsychol Soc. 2009;15(5):645–9. Epub 2009/07/28. doi: 10.1017/S1355617709990348. PubMed PMID: 19631024; PubMed Central PMCID: PMCPMC2791360.

128. Abel TJ, Rhone AE, Nourski KV, Kawasaki H, Oya H, Griffiths TD, et al. Direct physiologic evidence of a heteromodal convergence region for proper naming in human left anterior temporal lobe. J Neurosci. 2015;35(4):1513–20. doi: 10.1523/JNEUROSCI.3387-14.2015. PubMed PMID: 25632128; PubMed Central PMCID: PMCPMC4308598.

129. Patterson K, Nestor PJ, Rogers TT. Where do you know what you know? The representation of semantic knowledge in the human brain. Nat Rev Neurosci. 2007;8(12):976–87. Epub 2007/11/21. doi: 10.1038/nrn2277. PubMed PMID: 18026167.

130. Scott SK, Blank CC, Rosen S, Wise RJ. Identification of a pathway for intelligible speech in the left temporal lobe. Brain. 2000;123 Pt 12:2400–6. Epub 2000/12/02. doi: 10.1093/brain/123.12.2400. PubMed PMID: 11099443; PubMed Central PMCID: PMCPMC5630088.

131. Spitsyna G, Warren JE, Scott SK, Turkheimer FE, Wise RJ. Converging language streams in the human temporal lobe. J Neurosci. 2006;26(28):7328–36. Epub 2006/07/14. doi: 10.1523/JNEUROSCI.0559-06.2006. PubMed PMID: 16837579; PubMed Central PMCID: PMCPMC6674192.

132. Gorno-Tempini ML, Dronkers NF, Rankin KP, Ogar JM, Phengrasamy L, Rosen HJ, et al. Cognition and anatomy in three variants of primary progressive aphasia. Ann Neurol. 2004;55(3):335–46. Epub 2004/03/03. doi: 10.1002/ana.10825. PubMed PMID: 14991811; PubMed Central PMCID: PMCPMC2362399.

133. Mesulam MM. From sensation to cognition. Brain. 1998;121 (Pt 6):1013–52.

134. Rolls ET. The cingulate cortex and limbic systems for emotion, action, and memory. Brain Struct Funct. 2019;224(9):3001–18. Epub 2019/08/28. doi: 10.1007/s00429-019-01945-2. PubMed PMID: 31451898; PubMed Central PMCID: PMCPMC6875144.

135. Engel AK, Gerloff C, Hilgetag CC, Nolte G. Intrinsic coupling modes: multiscale interactions in ongoing brain activity. Neuron. 2013;80(4):867–86. Epub 2013/11/26. doi: 10.1016/j.neuron.2013.09.038. PubMed PMID: 24267648.

136. Ojemann GA, Ojemann J, Ramsey NF. Relation between functional magnetic resonance imaging (fMRI) and single neuron, local field potential (LFP) and electrocorticography (ECoG) activity in human cortex. Front Hum Neurosci. 2013;7:34. Epub 2013/02/23. doi: 10.3389/fnhum.2013.00034. PubMed PMID: 23431088; PubMed Central PMCID: PMCPMC3576621.

137. Makris N, Papadimitriou GM, Kaiser JR, Sorg S, Kennedy DN, Pandya DN. Delineation of the middle longitudinal fascicle in humans: a quantitative, in vivo, DT-MRI study. Cereb Cortex. 2009;19(4):777–85. Epub 2008/08/02. doi: 10.1093/cercor/bhn124. PubMed PMID: 18669591; PubMed Central PMCID: PMCPMC2651473.

138. Binney RJ, Parker GJ, Lambon Ralph MA. Convergent connectivity and graded specialization in the rostral human temporal lobe as revealed by diffusion-weighted imaging probabilistic tractography. J Cogn Neurosci. 2012;24(10):1998–2014. Epub 2012/06/23. doi: 10.1162/jocn_a_00263. PubMed PMID: 22721379.

139. Gonzalez Alam T, McKeown BLA, Gao Z, Bernhardt B, Vos de Wael R, Margulies DS, et al. A tale of two gradients: differences between the left and right hemispheres predict semantic cognition. Brain Struct Funct. 2021. Epub 2021/09/13. doi: 10.1007/s00429-021-02374-w. PubMed PMID: 34510282.

140. Dobbins IG, Wagner AD. Domain-general and domain-sensitive prefrontal mechanisms for recollecting events and detecting novelty. Cereb Cortex. 2005;15(11):1768–78. Epub 2005/02/25. doi: 10.1093/cercor/bhi054. PubMed PMID: 15728740.

141. Hartwigsen G, Bengio Y, Bzdok D. How does hemispheric specialization contribute to human-defining cognition? Neuron. 2021;109(13):2075–90. Epub 2021/05/19. doi: 10.1016/j.neuron.2021.04.024. PubMed PMID: 34004139; PubMed Central PMCID: PMCPMC8273110.

142. Sherman SM, Guillery RW. Distinct functions for direct and transthalamic corticocortical connections. J Neurophysiol. 2011;106(3):1068–77. doi: jn.00429.2011 [pii]; 10.1152/jn.00429.2011 [doi].

143. Hu B. Functional organization of lemniscal and nonlemniscal auditory thalamus. Exp Br Res. 2003;153(4):543–9.

144. Seitzman BA, Snyder AZ, Leuthardt EC, Shimony JS. The State of Resting State Networks. Top Magn Reson Imaging. 2019;28(4):189–96. Epub 2019/08/07. doi: 10.1097/RMR.0000000000000214. PubMed PMID: 31385898; PubMed Central PMCID: PMCPMC6686880.

145. Feinsinger A, Pouratian N, Ebadi H, Adolphs R, Andersen R, Beauchamp MS, et al. Ethical commitments, principles, and practices guiding intracranial neuroscientific research in humans. Neuron. 2022;110(2):188–94. Epub 2022/01/21. doi: 10.1016/j.neuron.2021.11.011. PubMed PMID: 35051364.

146. Nourski KV, Howard MA, 3rd. Invasive recordings in the human auditory cortex. Handb Clin Neurol. 2015;129:225–44. doi: 10.1016/B978-0-444-62630-1.00013-5. PubMed PMID: 25726272.

147. Jenkinson M, Bannister P, Brady M, Smith S. Improved optimization for the robust and accurate linear registration and motion correction of brain images. Neuroimage. 2002;17(2):825–41. Epub 2002/10/16. PubMed PMID: 12377157.

148. Rohr K, Stiehl HS, Sprengel R, Buzug TM, Weese J, Kuhn MH. Landmark-based elastic registration using approximating thin-plate splines. IEEE Trans Med Imaging. 2001;20(6):526–34. Epub 2001/07/05. doi: 10.1109/42.929618. PubMed PMID: 11437112.

149. Dale AM, Fischl B, Sereno MI. Cortical surface-based analysis. I. Segmentation and surface reconstruction. Neuroimage. 1999;9(2):179–94. Epub 1999/02/05. doi: 10.1006/nimg.1998.0395. PubMed PMID: 9931268.

150. Fischl B, Salat DH, Busa E, Albert M, Dieterich M, Haselgrove C, et al. Whole brain segmentation: automated labeling of neuroanatomical structures in the human brain. Neuron. 2002;33(3):341–55. Epub 2002/02/08. doi: 10.1016/s0896-6273(02)00569-x. PubMed PMID: 11832223.

151. Destrieux C, Fischl B, Dale A, Halgren E. Automatic parcellation of human cortical gyri and sulci using standard anatomical nomenclature. Neuroimage. 2010;53(1):1–15. doi: 10.1016/j.neuroimage.2010.06.010. PubMed PMID: 20547229; PubMed Central PMCID: PMCPMC2937159.

152. Destrieux C, Terrier LM, Andersson F, Love SA, Cottier JP, Duvernoy H, et al. A practical guide for the identification of major sulcogyral structures of the human cortex. Brain Struct Funct. 2017;222(4):2001–15. doi: 10.1007/s00429-016-1320-z. PubMed PMID: 27709299.

153. Jenkinson M, Beckmann CF, Behrens TE, Woolrich MW, Smith SM. FSL. Neuroimage. 2012;62(2):782-90. Epub 2011/1008. doi: 10.1016/j.neuroimage.2011.09.015. PubMed PMID: 21979382.

154. Brugge JF, Nourski KV, Oya H, Reale RA, Kawasaki H, Steinschneider M, et al. Coding of repetitive transients by auditory cortex on Heschl’s gyrus. J Neurophysiol. 2009;102(4):2358–74. doi: 10.1152/jn.91346.2008. PubMed PMID: 19675285; PubMed Central PMCID: PMCPMC2775384.

155. Nourski KV, Steinschneider M, Rhone AE. Electrocorticographic Activation within Human Auditory Cortex during Dialog-Based Language and Cognitive Testing. Front Hum Neurosci. 2016;10:202. doi: 10.3389/fnhum.2016.00202. PubMed PMID: 27199720; PubMed Central PMCID: PMCPMC4854871.

156. Power JD, Barnes KA, Snyder AZ, Schlaggar BL, Petersen SE. Spurious but systematic correlations in functional connectivity MRI networks arise from subject motion. Neuroimage. 2012;59(3):2142–54. Epub 2011/10/25. doi: 10.1016/j.neuroimage.2011.10.018. PubMed PMID: 22019881; PubMed Central PMCID: PMCPMC3254728.

157. Kovach CK, Gander PE. The demodulated band transform. J Neurosci Methods. 2016;261:135–54. doi: 10.1016/j.jneumeth.2015.12.004. PubMed PMID: 26711370; PubMed Central PMCID: PMCPMC5084918.

158. Satopaa V, Albrecht J, Irwin D, Raghavan B, editors. Finding a “Kneedle” in a Haystack: Detecting Knee Points in System Behavior. 2011 31st International Conference on Distributed Computing Systems Workshops; 2011 20-24 June 2011.

159. Field CA, Welsh AH. Bootstrapping clustered data. Journal of the Royal Statistical Society: Series B (Statistical Methodology). 2007;69(3):369–90. doi: https://doi.org/10.1111/j.1467-9868.2007.00593.x.

160. Ren S, Lai H, Tong W, Aminzadeh M, Hou X, Lai S. Nonparametric bootstrapping for hierarchical data. Journal of Applied Statistics. 2010;37(9):1487–98. doi: 10.1080/02664760903046102.

