## Supplementary Figures and Tables for "Functional geometry of auditory cortical resting state networks derived from intracranial electrophysiology"

**Supplementary Table 1.** Participant demographics and study information.

| Participant | Age | Sex | Handedness | Language  dominance | Subdural  sites | Depth sites | Lang. dom. dataset  arrays | fMRI dataset  arrays | Seizure focus |
| --- | --- | --- | --- | --- | --- | --- | --- | --- | --- |
| L275 | 30 | M | R | L | 193 | 17 | Y | N | L lateral ventral temporal, R posterior ventral frontal |
| L282 | 40 | M | R | ? | 0 | 23 | Y | N | L posterior superior temporal, L posterior lateral ventral frontal |
| L287 | 18 | M | R | ? | 77 | 16 | Y | N | L parietal |
| R288 | 21 | M | R | L | 164 | 26 | Y | N | R temporal pole |
| L292 | 50 | F | L | R | 95 | 49 | Y | N | L medial temporal |
| R294 | 35 | M | R | ? | 127 | 57 | Y | N | R dorsal frontal pole |
| L307 | 32 | M | R | L | 171 | 30 | Y | N | L posterior insula |
| R316 | 31 | F | R | ? | 71 | 20 | Y | N | R medial temporal |
| R320 | 51 | F | R | ? | 191 | 21 | Y | N | R medial temporal |
| R322 | 28 | F | R | ? | 48 | 30 | Y | N | R posterior medial frontal |
| R329 | 20 | M | R | ? | 75 | 36 | Y | N | R medial frontal |
| R334 | 39 | M | L | L | 183 | 25 | Y | N | R posterior medial temporal |
| R335 | 32 | M | R | L | 38 | 34 | Y | N | bilateral medial temporal |
| L357 | 36 | M | R | L | 99 | 39 | Y | N | L medial temporal |
| L362 | 59 | M | R | B | 89 | 33 | N | N | bilateral multiple locations |
| R369 | 30 | M | R | L | 172 | 44 | Y | Y | R medial temporal |
| L372 | 34 | M | R | L | 147 | 32 | Y | Y | L temporal pole |
| R376 | 48 | F | R | L | 164 | 42 | Y | Y | R medial temporal |
| R399 | 22 | F | R | L | 153 | 27 | Y | N | R temporal (uncertain medial & neocortical) |
| L400 | 59 | F | R | L | 111 | 38 | Y | Y | L amygdala (medial temporal) |
| L403 | 56 | F | R | L | 166 | 25 | Y | N | L medial temporal |
| L405 | 19 | M | R | L | 73 | 51 | Y | Y | L frontal (posterior lateral) |
| L409 | 31 | F | R | L | 143 | 17 | Y | Y | L medial temporal, L temporal pole |
| R413 | 22 | M | L | R | 177 | 41 | Y | N | R medial temporal |
| L416 | 33 | M | B | L | 61 | 36 | Y | N | L ventral occipital, L lateral temporal |
| R418 | 25 | F | R | L | 50 | 33 | Y | Y | R temporal (medial & lateral posterior cortex) |
| L423 | 51 | M | R | L | 116 | 37 | Y | Y | L medial temporal |
| L425 | 52 | M | L | ? | 153 | 35 | N | N | L medial temporal |
| R429 | 32 | F | R | L | 101 | 40 | Y | N | R medial temporal and temporal pole |
| R434 | 39 | F | R | L | 160 | 41 | Y | N | R medial temporal |
| L439 | 37 | M | R | L | 0 | 32 | Y | N | R medial frontal (posterior dorsal) |
| L442 | 33 | F | R | L | 146 | 20 | Y | N | multifocal L frontal, temporal, parietal |
| R456 | 31 | M | L | ? | 156 | 40 | N | Y | R medial temporal, R posterior lateral superior temporal |
| L457 | 18 | M | R | ? | 0 | 66 | Y | N | L medial temporal |
| R458 | 23 | M | R | L | 0 | 48 | Y | N | R temporal (middle & posterior lateral) |
| L460 | 52 | M | R | L | 113 | 41 | Y | Y | L hippocampus (medial temporal) |
| L477 | 23 | F | R | L | 94 | 23 | Y | N | L parietal |
| L514 | 46 | M | R | L | 0 | 117 | Y | N | L insula (anterior) |
| R515 | 21 | F | R | L | 0 | 125 | Y | N | R medial temporal |
| R524 | 18 | M | L | L | 0 | 83 | Y | N | R medial temporal, L medial temporal |
| L525 | 46 | F | R | L | 149 | 59 | Y | N | Bilateral multifocal (predominantly L frontal & temporal) |
| R532 | 42 | F | R | L | 134 | 46 | Y | N | R ventral frontal (posterior) |
| L538 | 18 | F | R | L | 0 | 109 | Y | N | L insula and orbitofrontal |
| R559 | 27 | F | R | L | 0 | 98 | Y | N | R inferior frontal (middle - posterior) |
| R567 | 33 | M | R | ? | 0 | 89 | Y | N | R insula |
| L585 | 39 | F | R | L | 99 | 41 | Y | N | L medial temporal |
| R610 | 21 | M | R | L | 0 | 97 | Y | N | R anterior cingulate |
| L625 | 24 | F | R | L | 0 | 82 | Y | N | L medial temporal |
| L634 | 22 | F | R | L | 0 | 72 | Y | N | L medial temporal |

**Supplementary Table 2.** ROIs and electrode coverage.

| **ROI group** | **ROI** | **ROI abbreviation** | **Overall average embedding** | | **Language dominance** | | **fMRI** | |
| --- | --- | --- | --- | --- | --- | --- | --- | --- |
|  |  |  | ***N*_participants_** | ***n*_sites_** | ***N*_participants_** | ***n*_sites_** | ***N*_participants_** | ***n*_sites_** |
| **Auditory core** | Heschl’s gyrus, posteromedial portion | HGPM | **38** | **175** | **36** | **164** | **10** | **49** |
| **Auditory non-core** | Heschl’s gyrus, anterolateral portion | HGAL | **30** | **111** | **29** | **106** | **7** | **22** |
|  | Planum temporale | PT | **21** | **63** | **19** | **58** | **8** | **22** |
|  | Planum polare | PP | **26** | **53** | **24** | **51** | **8** | **22** |
|  | Superior temporal gyrus, posterior portion | STGP | **37** | **355** | **34** | **315** | **10** | **83** |
|  | Superior temporal gyrus, middle portion | STGM | **36** | **225** | **33** | **193** | **10** | **53** |
| **Auditory-related** | Superior temporal gyrus, anterior portion | STGA | **29** | **65** | **27** | **61** | **8** | **18** |
|  | Posterior insula | InsP | **33** | **109** | **32** | **107** | **9** | **15** |
|  | Superior temporal sulcus, upper bank | STSU | **23** | **107** | *22* | *106* | *4* | *13* |
|  | Superior temporal sulcus, lower bank | STSL | **26** | **92** | **25** | **90** | **5** | **17** |
|  | Middle temporal gyrus, posterior portion | MTGP | **37** | **411** | **35** | **377** | **10** | **96** |
|  | Middle temporal gyrus, middle portion | MTGM | **39** | **284** | **37** | **255** | **10** | **72** |
|  | Middle temporal gyrus, anterior portion | MTGA | **33** | **114** | **31** | **103** | **9** | **34** |
|  | Supramarginal gyrus | SMG | **40** | **355** | **37** | **316** | **10** | **93** |
|  | Angular gyrus, posterior portion | AGP | **29** | **155** | *27* | *145* | **9** | **64** |
|  | Angular gyrus, anterior portion | AGA | **15** | **66** | **14** | **64** | **6** | **26** |
| **Prefrontal** | Inferior frontal gyrus, pars opercularis | IFGop | **35** | **109** | **32** | **100** | **10** | **32** |
|  | Inferior frontal gyrus, pars triangularis | IFGtr | **39** | **192** | **37** | **183** | **9** | **48** |
|  | Inferior frontal gyrus, pars orbitalis | IFGor | **20** | **45** | **18** | **40** | **4** | **9** |
|  | Middle frontal gyrus | MFG | **43** | **517** | **40** | **462** | **9** | **105** |
|  | Superior frontal gyrus | SFG | **25** | **211** | *23* | *191* | *5* | *59* |
|  | Anterior cingulate cortex | ACC | **26** | **44** | *23* | *41* | **3** | **4** |
|  | Orbital gyri | OG | **43** | **382** | **40** | **360** | **10** | **104** |
|  | Transverse frontopolar gyrus | TFG | **20** | **85** | *19* | *77* | **7** | **23** |
|  | Frontomarginal gyrus | FMG | *4* | *9* | *4* | *9* | *0* | *0* |
| **Sensorimotor** | Precentral gyrus, dorsal portion | dPreCG | **29** | **106** | **27** | **101** | **7** | **19** |
|  | Postcentral gyrus, dorsal portion | dPostCG | **25** | **93** | *23* | *89* | **4** | **13** |
|  | Precentral gyrus, ventral portion | vPreCG | **42** | **237** | **39** | **220** | **10** | **48** |
|  | Postcentral gyrus, ventral portion | vPostCG | **38** | **190** | **35** | **173** | **9** | **47** |
|  | Paracentral lobule | ParaCL | *7* | *15* | *7* | *15* | *0* | *0* |
| **Other** | Premotor cortex | PMC | **41** | **175** | **38** | **162** | **9** | **33** |
|  | Cingulate gyrus, middle portion | CingM | **20** | **37** | *17* | *33* | **4** | **7** |
|  | Cingulate gyrus, posterior portion / Precuneus | PCC/pC | **21** | **84** | *19* | *79* | *2* | *5* |
|  | Parahippocampal gyrus | PHG | **28** | **109** | **26** | **105** | **8** | **22** |
|  | Fusiform gyrus | FG | **33** | **152** | **31** | **147** | **9** | **24** |
|  | Inferior temporal gyrus, posterior portion | ITGP | **24** | **85** | **23** | **83** | **8** | **23** |
|  | Inferior temporal gyrus, middle portion | ITGM | **31** | **111** | **29** | **108** | **7** | **22** |
|  | Inferior temporal gyrus, anterior portion | ITGA | **28** | **110** | **26** | **102** | **8** | **36** |
|  | Temporal pole | TP | **33** | **280** | **30** | **252** | **10** | **74** |
|  | Anterior insula | InsA | **29** | **77** | **27** | **74** | **9** | **16** |
|  | Frontal operculum | fOperc | **14** | **23** | *12* | *19* | *4* | *7* |
|  | Parietal operculum | pOperc | *7* | *21* | *7* | *21* | *1* | *2* |
|  | Gyrus rectus | GR | **33** | **66** | **31** | **58** | **9** | **15** |
|  | Subcallosal gyrus | SubcG | *5* | *8* | *3* | *6* | *1* | *1* |
|  | Superior parietal lobule / Intraparietal sulcus | SPL | **16** | **92** | **14** | **82** | **4** | **13** |
|  | Cuneus | Cun | *4* | *7* | *4* | *7* | *2* | *3* |
|  | Lingual gyrus | LingG | **10** | **24** | *10* | *24* | *3* | *5* |
|  | Occipital pole | OP | *2* | *4* | *2* | *4* | *0* | *0* |
|  | Superior occipital gyrus | SOG | *4* | *9* | *4* | *9* | *0* | *0* |
|  | Middle occipital gyrus | MOG | **18** | **73** | *18* | *73* | **4** | **21** |
|  | Inferior occipital gyrus | IOG | *8* | *20* | *8* | *20* | *2* | *7* |
|  | Amygdala | Amyg | **32** | **80** | **30** | **78** | *6* | *19* |
|  | Hippocampus | Hipp | **31** | **87** | **30** | **85** | **5** | **14** |
|  | Putamen | Put | *9* | *15* | *9* | *15* | *3* | *4* |
|  | Globus pallidus | GP | *1* | *1* | *0* | *0* | *1* | *1* |
|  | Caudate nucleus | Caud | *5* | *10* | *5* | *10* | *0* | *0* |
|  | Substantia innominata | SubInn | *2* | *5* | *2* | *5* | *2* | *5* |
|  | Ventral striatum | vStr | *1* | *2* | *1* | *2* | *1* | *2* |
| **Total** | | | 49 | 6742 | 46 | 6235 | 10 | 1591 |

Participant and site numbers that were not included in group averages are denoted in *gray italics*.

**Supplementary Table 3.** List of abbreviations.

| ACC | Anterior cingulate cortex |
| --- | --- |
| AGA | Angular gyrus, anterior portion |
| AGP | Angular gyrus, posterior portion |
| Amyg | Amygdala |
| ATL | Anterior temporal lobe |
| Caud | Caudate nucleus |
| CingM | Cingulate gyrus, middle portion |
| Cun | Cuneus |
| DME | Diffusion map embedding |
| dPostCG | Postcentral gyrus, dorsal portion |
| dPreCG | Precentral gyrus, dorsal portion |
| FG | Fusiform gyrus |
| FMG | Frontomarginal gyrus |
| fMRI | Functional magnetic resonance imaging |
| fOperc | Frontal operculum |
| GP | Globus pallidus |
| GR | Gyrus rectus |
| HGAL | Heschl’s gyrus, anterolateral portion |
| HGPM | Heschl’s gyrus, posteromedial portion |
| Hipp | Hippocampus |
| iEEG | Intracranial electroencephalography |
| IFGop | Inferior frontal gyrus, pars opercularis |
| IFGor | Inferior frontal gyrus, pars orbitalis |
| IFGtr | Inferior frontal gyrus, pars triangularis |
| InsA | Anterior insula |
| InsP | Posterior insula |
| IOG | Inferior occipital gyrus |
| ITGA | Inferior temporal gyrus, anterior portion |
| ITGM | Inferior temporal gyrus, middle portion |
| ITGP | Inferior temporal gyrus, posterior portion |
| LingG | Lingual gyrus |
| MFG | Middle frontal gyrus |
| MNI | Montreal Neurological Institute |
| MOG | Middle occipital gyrus |
| MRI | Magnetic resonance imaging |
| MTGA | Middle temporal gyrus, anterior portion |
| MTGM | Middle temporal gyrus, middle portion |
| MTGP | Middle temporal gyrus, posterior portion |
| OG | Orbital gyri |
| OP | Occipital pole |
| ParaCL | Paracentral lobule |
| PCC/pC | Cingulate gyrus, posterior portion / Precuneus |
| PHG | Parahippocampal gyrus |
| PMC | Premotor cortex |
| pOperc | Parietal operculum |
| PP | Planum polare |
| PT | Planum temporale |
| Put | Putamen |
| ROI | Region of interest |
| RS | Resting state |
| SFG | Superior frontal gyrus |

**Supplementary Table 3.** List of abbreviations *(continued)*.

| SMG | Supramarginal gyrus |
| --- | --- |
| SNR | Signal to noise ratio |
| SOG | Superior occipital gyrus |
| SPL | Superior parietal lobule / Intraparietal sulcus |
| STGA | Superior temporal gyrus, anterior portion |
| STGM | Superior temporal gyrus, middle portion |
| STGP | Superior temporal gyrus, posterior portion |
| STSL | Superior temporal sulcus, lower bank |
| STSU | Superior temporal sulcus, upper bank |
| SubcG | Subcallosal gyrus |
| SubInn | Substantia innominata |
| TFG | Transverse frontopolar gyrus |
| TP | Temporal pole |
| vPostCG | Postcentral gyrus, ventral portion |
| vPreCG | Precentral gyrus, ventral portion |
| vStr | Ventral striatum |

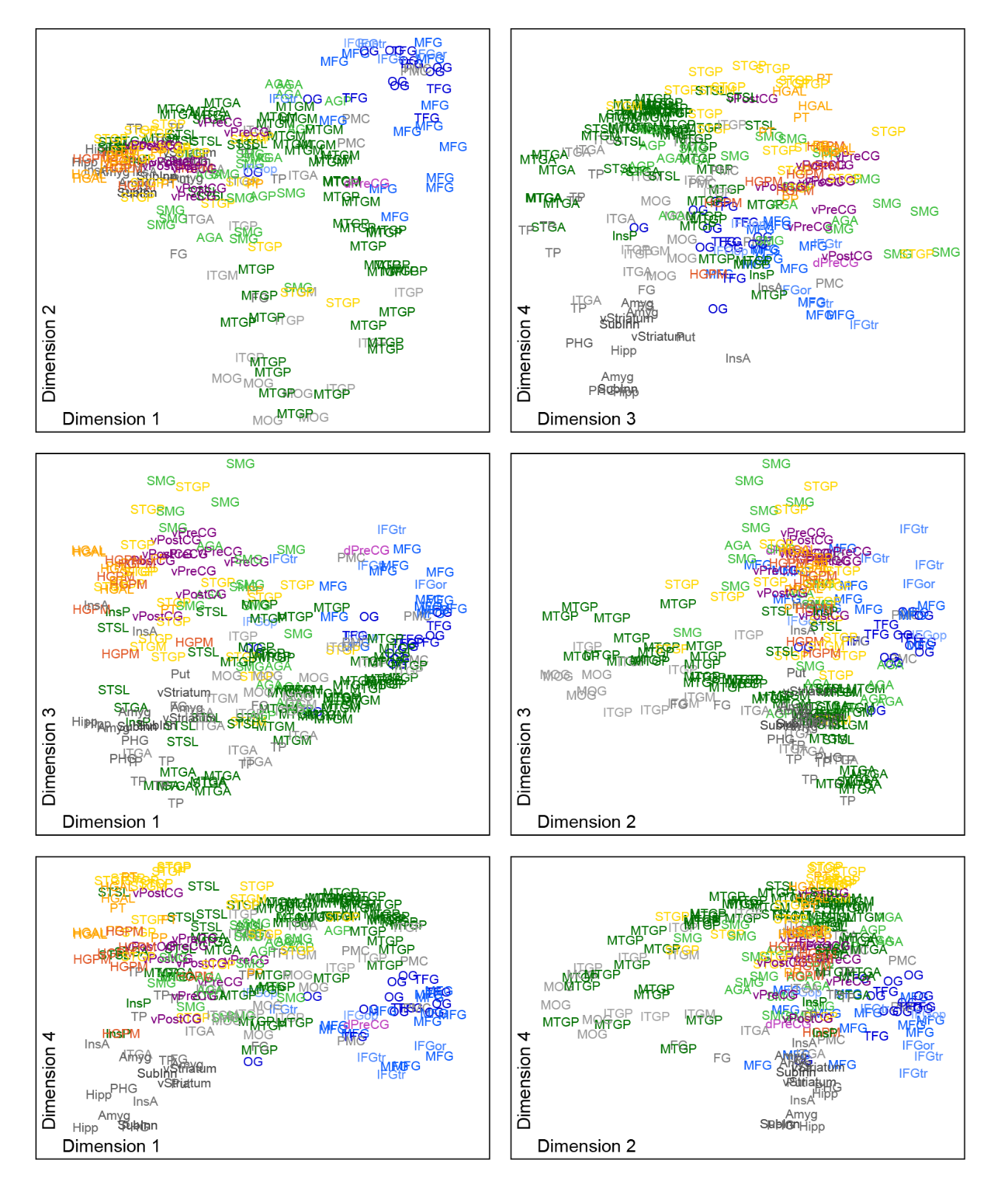

**Supplementary Figure 1**. Embedding plots in participant R376 plotted on the same scale in four dimensions.

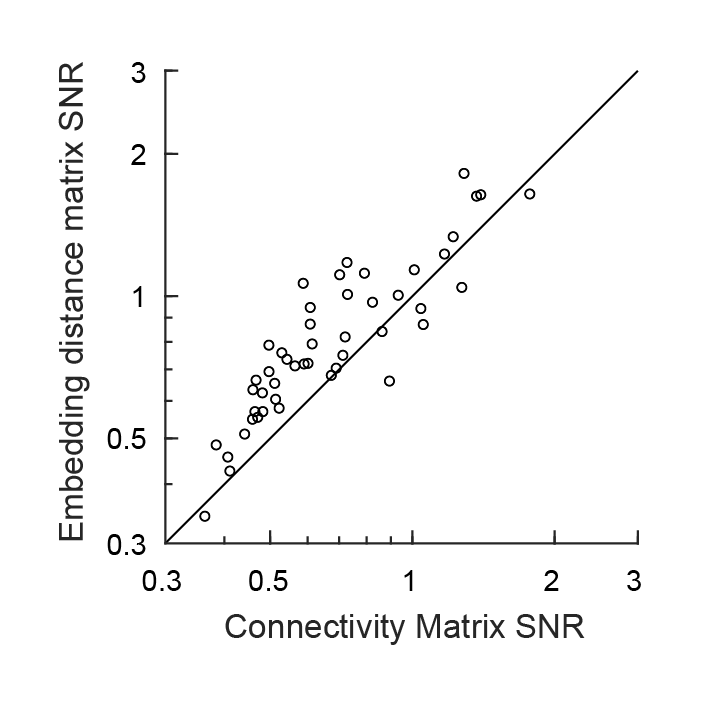

**Supplementary Figure 2**. Comparison of signal to noise ratio (SNR) for the embedding analysis versus direct analysis of functional connectivity. Each symbol corresponds to one participant. For each participant, the SNRs of embedding distances and connectivity were calculated from the recorded RS block as described in Methods. In most participants, the embedding analysis exhibited superior SNR characteristics compared to direct analysis of connectivity.

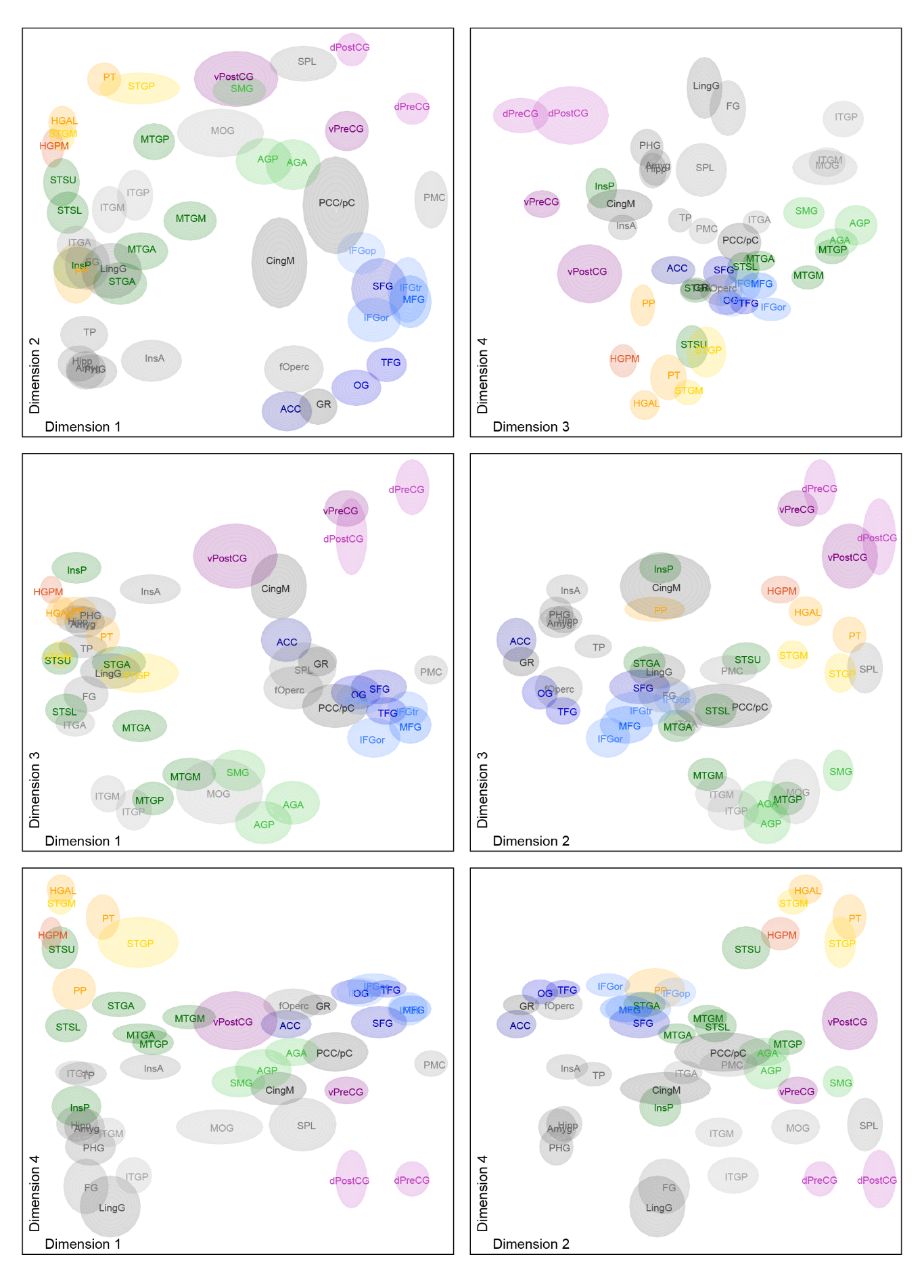

**Supplementary Figure 3.** Average gamma-band embedding, plotted on the same scale in the first four dimensions.

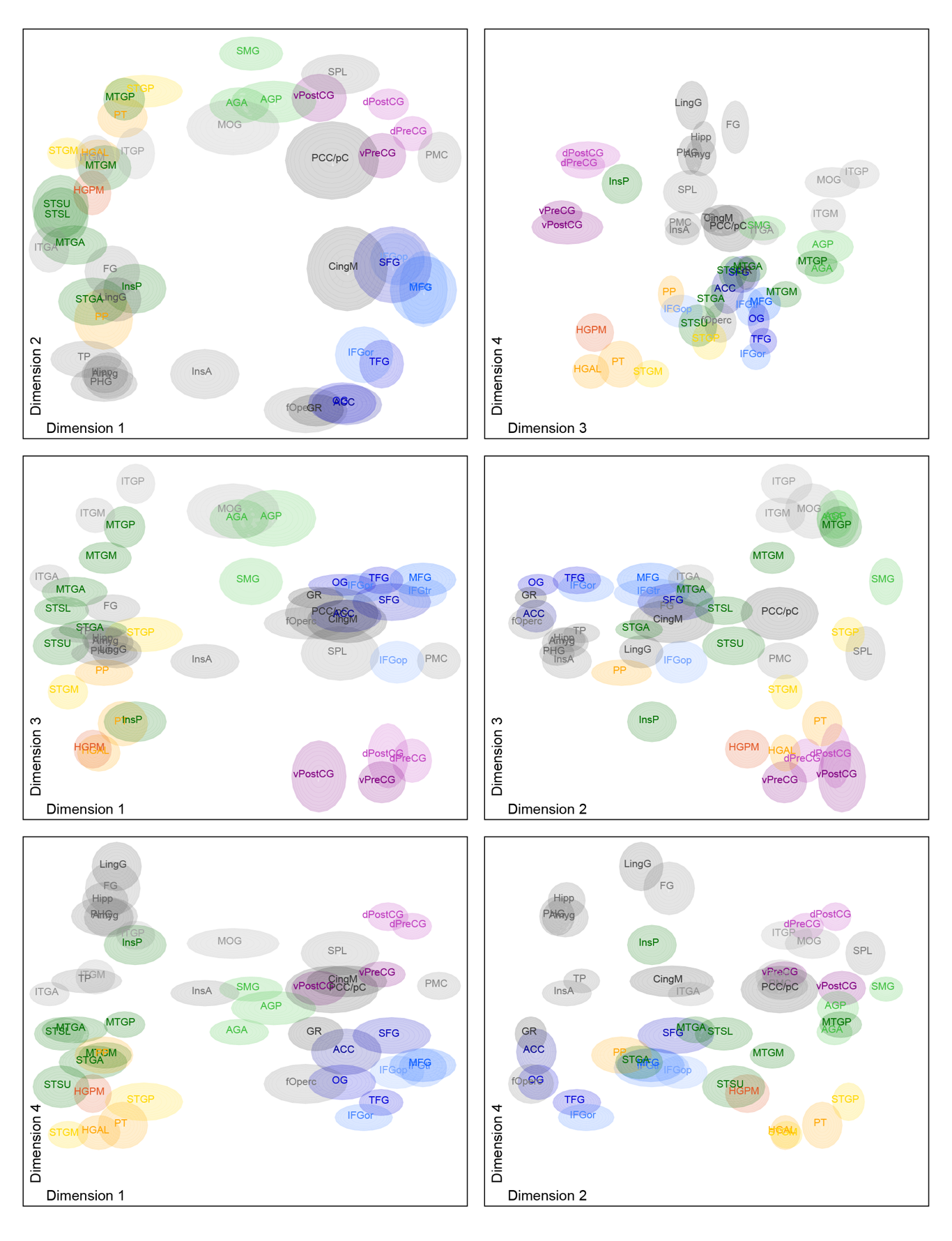
**Supplementary Figure 4.** Average beta-band embedding, plotted on the same scale in the first four dimensions

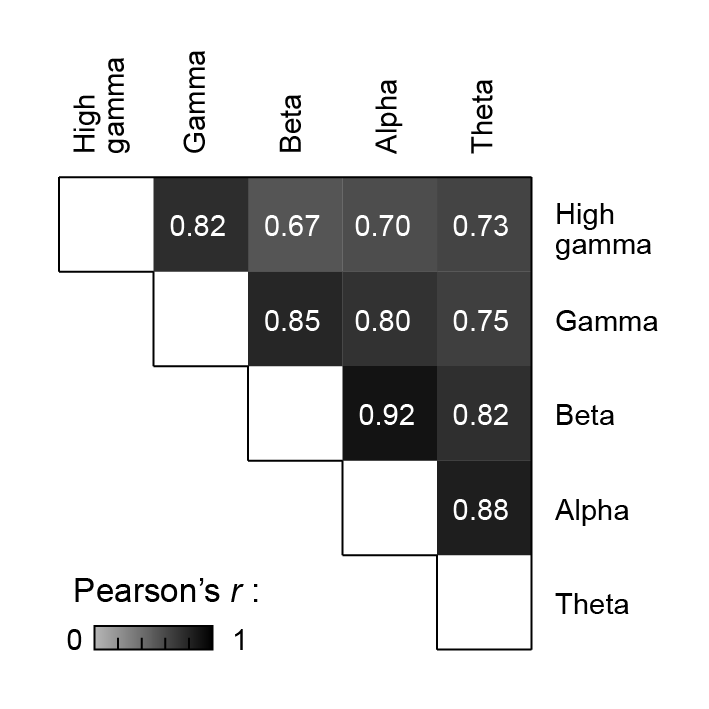

**Supplementary Figure 5.** Comparison of embedding results across frequency bands and functional connectivity measures. Data shown are Pearson correlations of embedding distances between measures and bands.

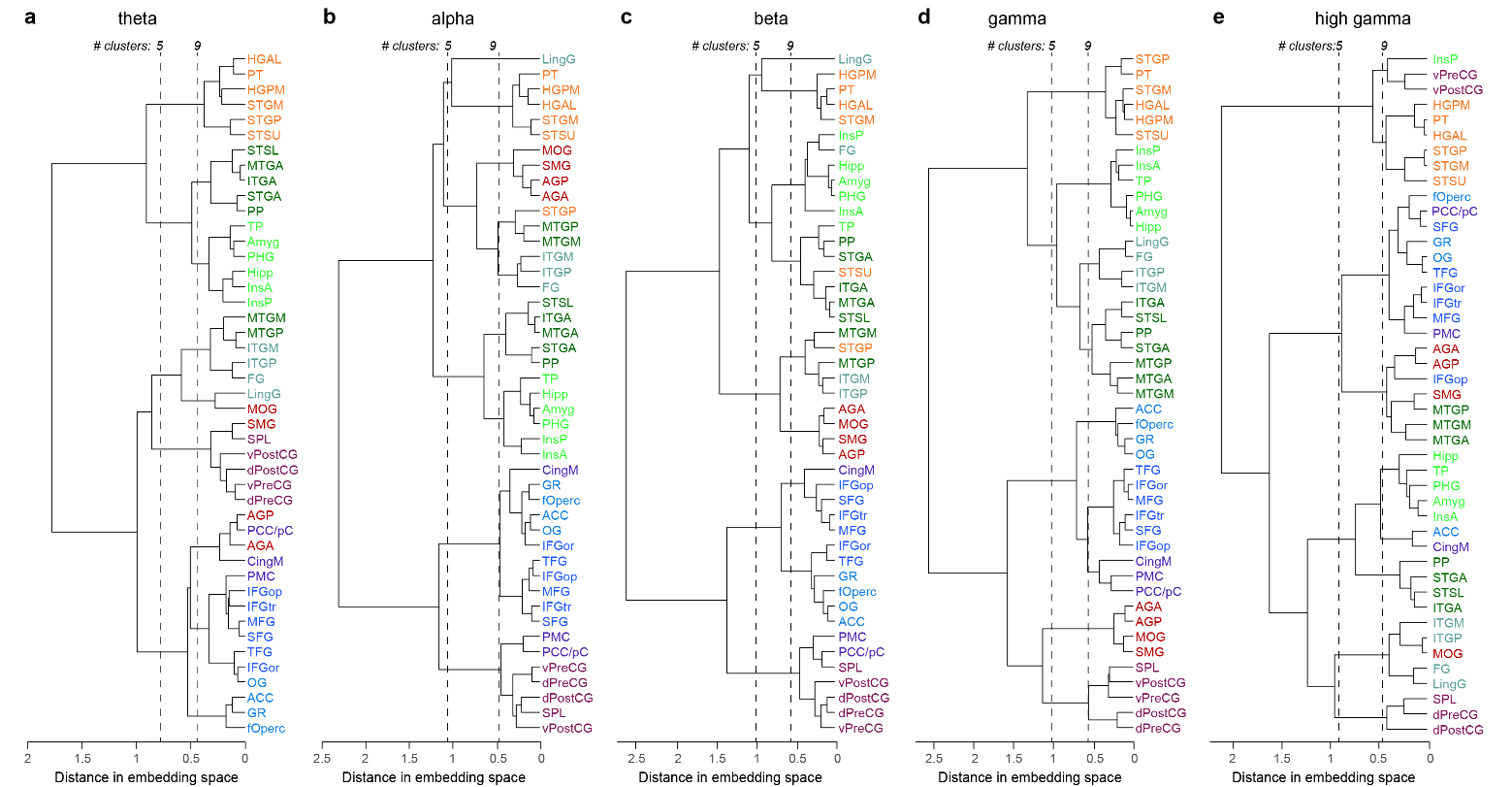

**Supplementary Figure 6.** Hierarchical clustering of data in embedding space for all studied bands. **a:** theta; **b:** alpha; **c:** beta; **d:** gamma (same data as in Figure 4); **e:** high gamma. Linkages between ROI groups identified using agglomerative clustering. As in Figure 4, two thresholds are shown for each band, n_Cluster_ = 5 and 9 (vertical dashed lines). The number of clusters is the number of lines in the dendrogram intersected by the threshold line. Clusters consist of all ROIs descending from the intersected line. The color scheme for ROI labels is set by the gamma parcellation.

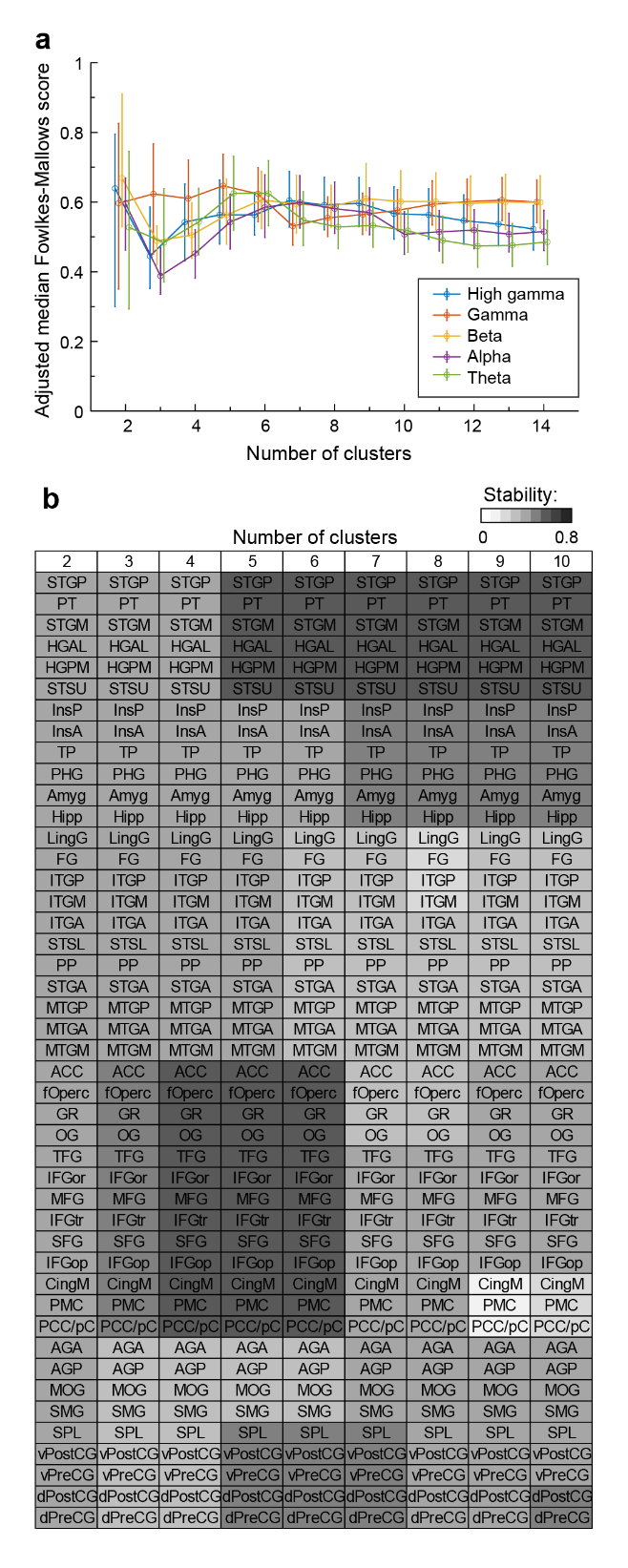

**Supplementary Figure 7. Stability of cluster results. a:** Stability of overall cluster results shown in Figure 4a and Supplementary Figure 6 was evaluated for each frequency band as a function cluster number using the Fowlkes-Mallows score. **b:** Cluster-wise stability for gamma band data as a function of cluster number was evaluated using the Jaccard index.

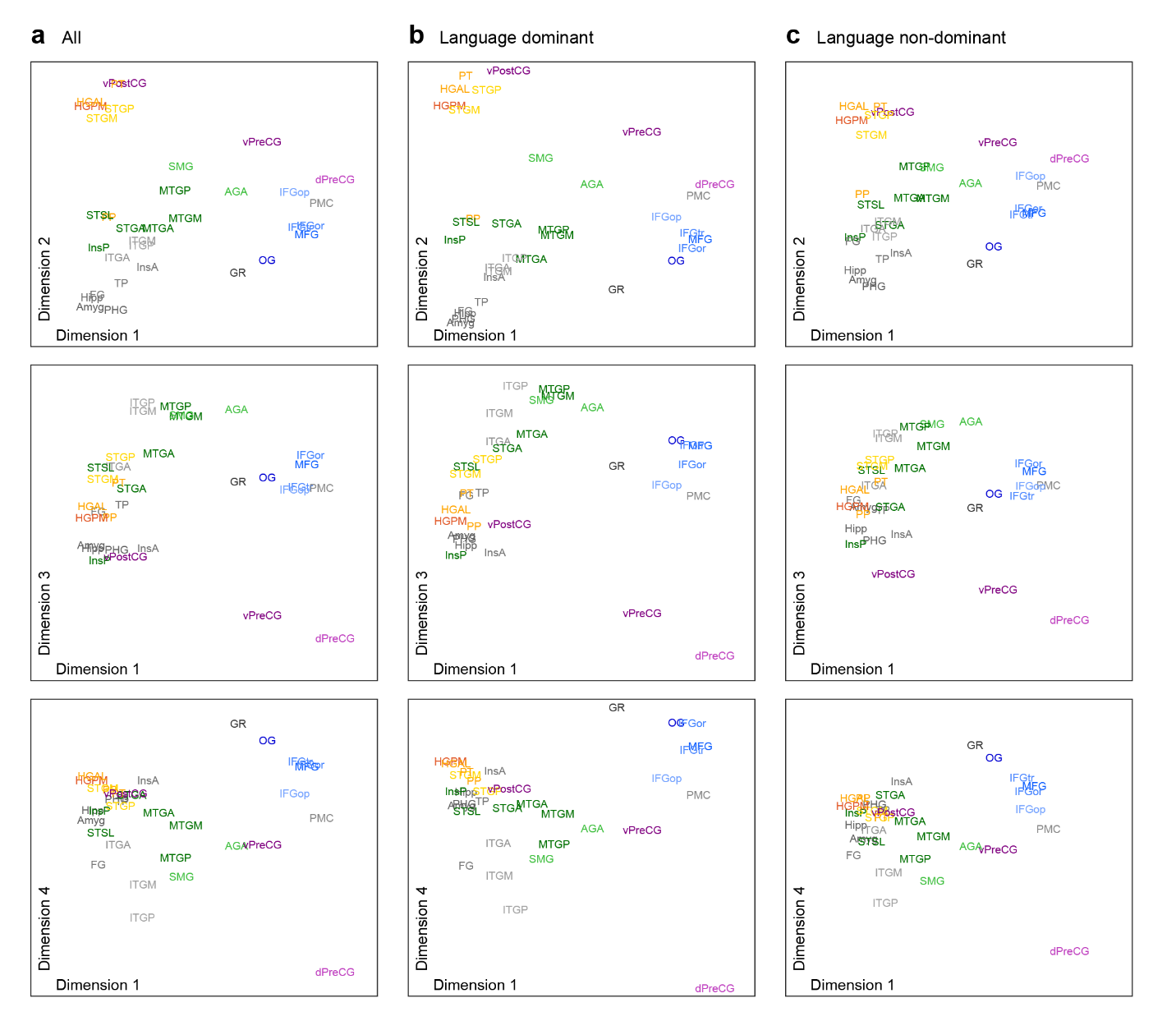

**Supplementary Figure 8. Auditory networks do not differ between hemispheres.** Data plotted on the same scale in the first 4 dimensions of embedding space for all dominant and non-dominant participants (**a**), just dominant (**b**), and just non-dominant (**c**).

**Supplementary Movie 1.** Average embedding data, visualized on the same scale in the first three dimensions. Same data as in Figure 3 and Supplementary Figure 3.

**Supplementary Movie 2.** Average embedding data, visualized on the same scale in the 2nd, 3rd, and 4th dimensions. Same data as in Figure 3 and Supplementary Figure 3.
